# Developmentally distinct architectures in top-down circuits

**DOI:** 10.1101/2023.08.27.555010

**Authors:** Cassandra B. Klune, Caitlin M. Goodpaster, Michael W. Gongwer, Christopher J. Gabriel, Rita Chen, Nico S. Jones, Lindsay A. Schwarz, Laura A. DeNardo

## Abstract

The medial prefrontal cortex (mPFC) plays a key role in learning, mood and decision making, including in how individuals respond to threats^1–6^. mPFC undergoes a uniquely protracted development, with changes in synapse density, cortical thickness, long-range connectivity, and neuronal encoding properties continuing into early adulthood^7–21^. Models suggest that before adulthood, the slow-developing mPFC cannot adequately regulate activity in faster-developing subcortical centers^22,23^. They propose that during development, the enhanced influence of subcortical systems underlies distinctive behavioural strategies of juveniles and adolescents and that increasing mPFC control over subcortical structures eventually allows adult behaviours to emerge. Yet it has remained unclear how a progressive strengthening of top-down control can lead to nonlinear changes in behaviour as individuals mature^24,25^. To address this discrepancy, here we monitored and manipulated activity in the developing brain as animals responded to threats, establishing direct causal links between frontolimbic circuit activity and the behavioural strategies of juvenile, adolescent and adult mice. Rather than a linear strengthening of mPFC synaptic connectivity progressively regulating behaviour, we uncovered multiple developmental switches in the behavioural roles of mPFC circuits targeting the basolateral amygdala (BLA) and nucleus accumbens (NAc). We show these changes are accompanied by axonal pruning coinciding with functional strengthening of synaptic connectivity in the mPFC-BLA and mPFC-NAc pathways, which mature at different rates. Our results reveal how developing mPFC circuits pass through distinct architectures that may make them optimally adapted to the demands of age-specific challenges.

## Main

Dispersal from parental care requires reorganization of behaviour, with prioritization of exploration and trial-and-error learning that eventually affords independence. Increasing demands of risky exploration likely put pressure on the brain systems that control learning, memory and decision making with respect to threatening stimuli^26^. But how brain development programs behavioural transitions remains poorly understood.

It is well established that in adults, mPFC circuits encode threatening cues and guide behavioural responses to threats through top-down control of subcortical regions including the BLA and NAc^27–41^. During development, the protracted maturation of mPFC coincides with conserved transitions in threat-induced behaviours and risk-taking^20,42,42–50^. In many species, while juveniles rely on caregivers for protection, adolescents engage in risky exploration and adults seek safety^24,42,44,45,47–50^. Prominent theories of brain and behavioural development have proposed that the slow maturation of mPFC initially allows subcortical emotion systems to dominate behavior, leading to enhanced impulsivity and risk taking in young individuals, particularly during adolescence^22,23,51^. Yet difficulties in explaining how progressive strengthening of top-down control can lead to nonlinear changes in behaviour and cognition have called some of these theories into question^24,25^.

To address these discrepancies, we developed a behavioural assay that allowed us to model conserved developmental changes in threat avoidance while enabling in-depth neuronal circuit interrogation. We show that compared to adults, juvenile and adolescent mice have lower levels of threat avoidance, with unique behavioural features at each age. With fibre photometry, we show that in the prelimbic cortex (PL), a subregion of mPFC that plays a key role in conditioned fear and threat avoidance^28,29,31,34,37,39,41^, in BLA, and in NAc, the neural dynamics underlying threat cues and threat avoidance behaviour mature along distinct trajectories. Using causal manipulations, viral circuit mapping and synaptic physiology, we show that PL circuits have distinct functional organization in different developmental windows. In juveniles, PL top-down circuits make weak, promiscuous synaptic connections in BLA and NAc that preclude mPFC’s role in threat-avoidance behaviour. In adolescents, rather than playing a passive role, activity in both the PL-BLA and PL-NAc pathways drives increased exploration at the expense of threat avoidance. Underlying these changes, synaptic excitation increases in the adolescent PL-NAc pathway while PL-BLA functions remain immature. These PL connections are ultimately refined in adults to establish high levels of threat avoidance and a circuit organization wherein activity in the PL-BLA pathway promotes threat avoidance and PL-NAc activity opposes it. Together, these findings reveal how the delicate coordination of synaptic maturation in parallel top-down pathways produce distinct mPFC circuit architectures that promote developmentally appropriate behaviours.

### Distinct threat avoidance behavioural strategies in juvenile, adolescent and adult mice

Mice exhibit developmental changes in the retention and expression of conditioned fear including infantile amnesia and a temporary suppression of conditioned fear in early adolescence^42,43,46,49,52^, but it is unclear to what extent threat avoidance behaviour changes as animals mature. To examine this, we refined the platform mediated avoidance assay^27^ for study of developing mice (PMA; Fig. 1a–c). In previous studies that used PMA, well-trained rats lever pressed for food until they heard a shock-conditioned tone that prompted them to enter a safety platform^27,28,53^. Because this assay required multiple days of food deprivation and operant training, it was incompatible with developmental studies. We refined PMA for study of developing mice by eliminating the food deprivation and lever press components and instead examining the balance between exploration and threat avoidance. To capture specific developmental timepoints, we limit training to a single day. To encourage exploration, we place sweet smelling odor pots underneath the shock floor where they are just out of reach. By analyzing keypoint tracking data with open-source software we previously developed, we perform high throughput, detailed, quantitative analysis of behavior across development.

**Figure 1.**
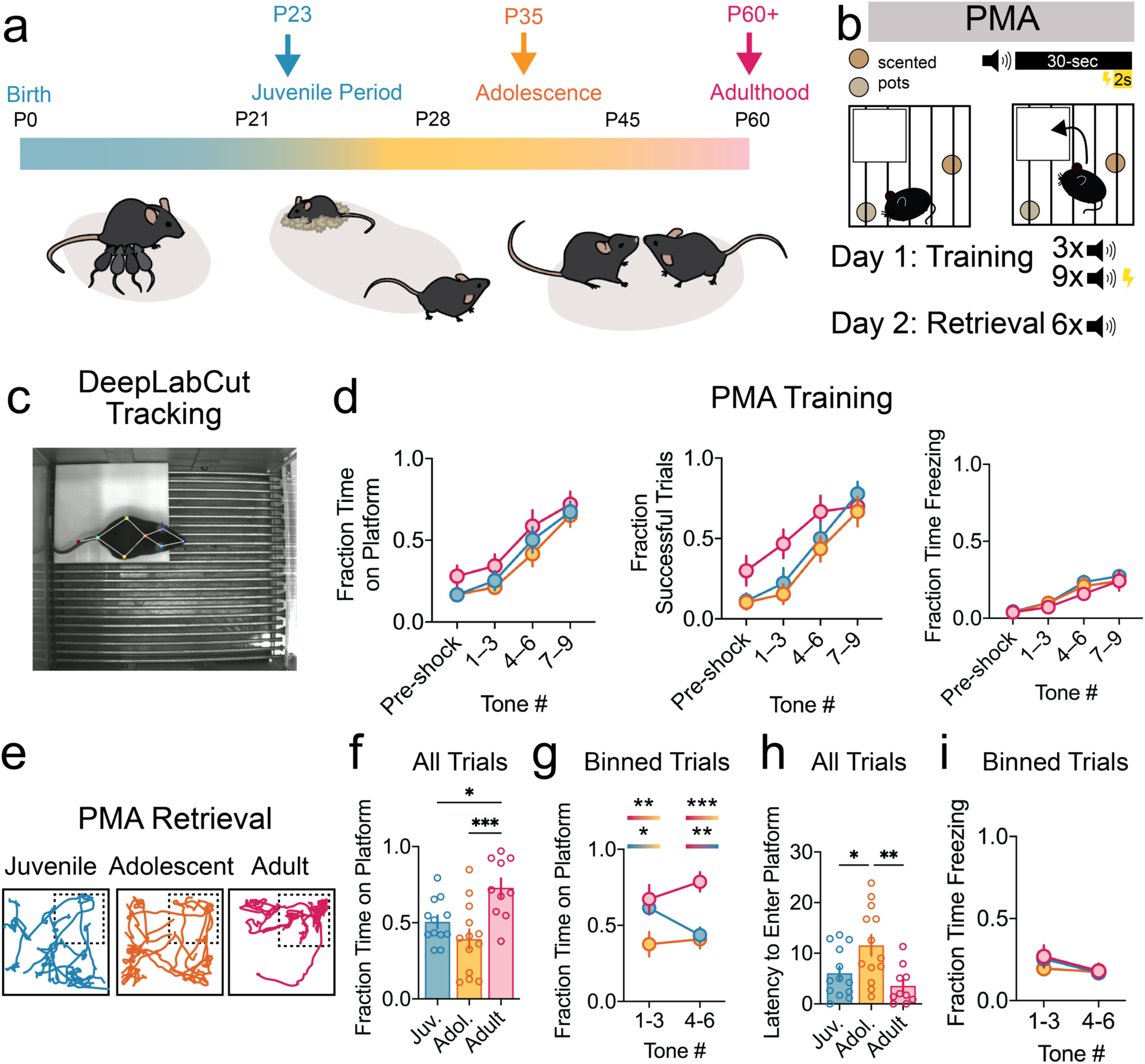
Learned avoidance behaviour is developmentally regulated. **a,** Selection of ages for study. **b,** PMA protocol. **c,** Keypoint tracking using DeepLabCut. **d,** Behavioural performance during PMA training (Juvenile: n=12, Adolescent: n=13, Adult: n=10; 2-way ANOVA). **e,** Representative examples of mouse trajectory maps during a PMA retrieval session. **f–i,** Behavioural performance during PMA retrieval (Juvenile: n=12, Adolescent: n=13, Adult: n=10). **f,** Time on the platform averaged across all tones (One-way ANOVA). **g,** Time on Platform averaged across bins of 3 tones (2-way ANOVA). **h,** Latency to enter platform averaged across 6 tones (One-way ANOVA). **i,** Freezing averaged across bins of 3 tones (2-way ANOVA). Data represent mean ± standard error of the mean (s.e.m.), *p<0.05, **p<0.01, ***p<0.001. Full statistical details can be found in Data Table 1.

Ages were chosen to span developmental milestones including weaning (postnatal day (P) 23), adolescence (P35) and adulthood (P60 and above). We did not observe age-dependent differences in the ability to learn PMA. Mice in all age groups similarly increased their fraction of successful trials and time spent on the safety platform and modestly increased conditioned freezing over the course of training (Fig. 1d). The next day, we tested their learned avoidance behaviour in a retrieval session. We presented conditioned tones in the absence of foot shocks and observed age-specific behavioural repertoires (Fig. 1e). During presentations of threatening tones, juveniles and adolescents spent less time on the platform compared to adults (Fig. 1f). To determine if the age groups organized their behavior differently over time, we compared time on the platform and conditioned freezing at the beginning versus the end of the retrieval session. Adults had persistent, high levels of PMA, adolescents had consistently low levels of PMA, and juveniles decreased PMA over the course of the retrieval session (Fig. 1g). Adolescents had the longest latency to enter the safety platform after the onset of the conditioned tone (Fig. 1h), consistent with elevated risky exploration at that age. Conditioned freezing levels were similar across ages (Fig. 1i), suggesting that developmental differences in PMA reflect differences in conditioned avoidance responses rather than in the strength of the association between the tone and the shock.

We next performed a series of control experiments to confirm that behavioural differences in PMA reflected developmental changes in learned threat avoidance behaviour. Shock sensitivity levels and distance traveled in the conditioning chamber were similar across age groups (Extended Data Fig. 1a,b). Adults spent significantly more time exploring the center of an open field compared to younger mice, suggesting that their robust PMA is not due to increased anxiety-like behaviour (Extended Data Fig. 1c). For each age group, non-shocked control mice spent significantly less time on the platform and had fewer ‘successful trials’, meaning they did not tend to be on the platform in the last two seconds on the tone when the shock would occur during training (Extended Data Fig. 2a–f). To determine if age-dependent changes in PMA during retrieval reflected developmental differences in the rate of fear extinction, in a cohort of mice, we performed a second retrieval test 22 days after training. We observed no significant changes in threat avoidance levels at any age, suggesting that fear extinction was not likely to determine the low levels of PMA observed in adolescents (Extended Data Fig. 3a–c). Together, these data suggest that observed differences in PMA reflect developmental changes in learned threat responses rather than changes in shock sensitivity, locomotion, innate anxiety levels, natural preference for being on the platform, or fear extinction.

### Region- and age-specific neural dynamics underlying PMA across development

In humans and rodents, PL, BLA, and NAc are key neural substrates for threat responding and learned threat avoidance^23,27,28,34–36,54–59^. Human studies have revealed that compared to children and adults, adolescent NAc and BLA have enhanced activation to emotional stimuli while PFC responses remain immature through adolescence^60,61^. In rodents, individual PL neurons encode threat-predictive cues, and conditioned freezing and PL activity is required for active threat avoidance^27,28,30,31,34,35,37,41,57^. BLA encodes learned threat associations and is required for the expression of active avoidance behaviour^5,6,27,62^. NAc promotes reward seeking behaviours and gates the expression of threat avoidance behaviour through its outputs to midbrain centers^56,63–65^. But how these brain regions respond during threat-predictive cues or threat-induced behaviours across development remains poorly understood^66^. To test this, we used fibre photometry to measure bulk calcium fluorescence as a proxy for neural activity during PMA retrieval in juvenile, adolescent, and adult mice (Fig 2a, b). Following PMA recordings viral expression and fiber placement were confirmed in all animals (Extended Data Fig 4a-c).

**Figure 2.**
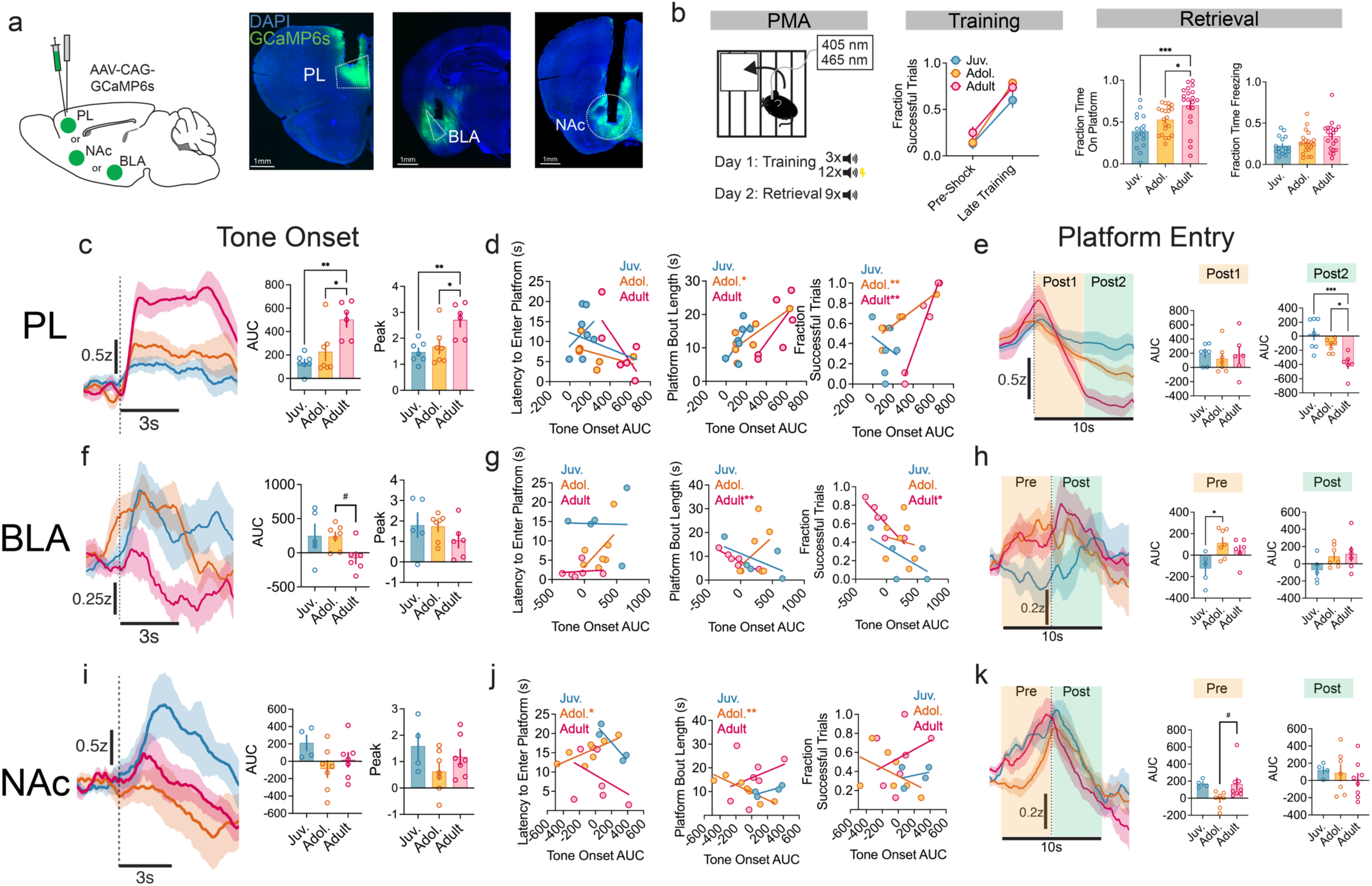
Neural dynamics underlying PMA in PL, BLA and NAc. **a,** Left: Schematic of AAV injection and optic fibre implant in PL, NAc or BLA. Right: Representative image showing GCaMP6s expression in PL, NAc and BLA neurons and fibre placement. **b,** Left: Schematic of PMA protocol. Right: Behavioural performance during PMA (Juvenile n=16, Adolescent n=21, Adult n=20, Successful trials: 2-way ANOVA; Time on platform and freezing, One-way ANOVA). **c-e**, Calcium recordings of PL during PMA retrieval (Juvenile n=7; Adolescent n=7, Adult n=6). **c,** Averaged Z-score in response to tone onset across the retrieval session (average only includes trials when the animal was off of the platform upon tone onset). Quantification of area under the curve (AUC) and Peak Z-score 3 seconds after tone onset (AUC: One-way ANOVA; Peak: One-way ANOVA). **d,** Correlations between the AUC during 3 seconds following tone onset and behaviour (Linear Regression). **e,** Averaged fluorescence changes (Z-score) surrounding platform entries (dotted line) across the entire retrieval session (9 tones total). Quantification of area under the curve (AUC) 0-5 seconds (Post1) and 5-10 seconds (Post2) after entry across ages (Post1: One-way ANOVA; Post2: One-way ANOVA). **f-h**, Calcium recordings of BLA during PMA retrieval (Juvenile n=5; Adolescent n=7, Adult n=6). **f,** Averaged Z-score in response to tone onset across the retrieval session (average only includes trials when the animal was off of the platform upon tone onset). Quantification of AUC and Peak Z-score value 3 seconds after tone onset (AUC: One-way ANOVA; Peak: One-way ANOVA). **g,** Correlations between the AUC during 3 seconds following tone onset and behaviour (Linear regression). **h,** Averaged Z-score surrounding platform entries (dotted line) across the retrieval session. Quantification of AUC-5 to 0 seconds before (Pre) and 0 to 5 seconds after (Post) entry across ages (Pre: One-way ANOVA; Post: One-way ANOVA). **i-k,** Calcium recordings of NAc during PMA retrieval (Juvenile n=4; Adolescent n=7, Adult n=7). **i,** Averaged Z-score in response to tone onset across the retrieval session (average only includes trials when the animal was off of the platform upon tone onset). Quantification of AUC and Peak Z-score value 3 seconds after tone onset (AUC: One-way ANOVA; Peak (One-way ANOVA). **j,** Correlations between the AUC during 3 seconds following tone onset and behavior across ages (Linear regression). **k,** Averaged Z-score surrounding platform entries (dotted line) across the retrieval session. Quantification of the total AUC-5 to 0 seconds (Pre) before entry to 0 to 5 seconds after (Post) entry across ages. **l,** Scale bars, 1 mm. Data represent mean ± s.e.m and shading reflects between-subjects s.e.m., ^#^p<0.10, *p<0.05, **p<0.01, ***p<0.001. Full statistical details can be found in Data Table 2.

During PMA retrieval, adults had significantly higher PL activity at tone onset compared to juveniles and adolescents (Fig. 2c). The area under the curve (AUC) of the tone response was positively correlated with behavioral measures in adolescents and adults but not in juveniles. In adolescents, the tone-evoked AUC was correlated with the platform bout duration (how long mice tended to stay on the platform before leaving again). In both adolescents and adults, the tone AUC was correlated with the fraction of successful trials (Fig. 2d). Mice of all ages had similar increases in PL activity during platform entries (Fig. 2e). However, once animals entered the safety platform, PL activity decreased to a greater extent in adults compared to juveniles and adolescents (Fig. 2e).

In the BLA, adults had low activity at tone onset, while juveniles and adolescents had moderate tone responses (Fig. 2f). The AUC of the tone onset response was negatively correlated with platform bout length and successful trials only in adults (Fig. 2g), indicating that low levels of BLA activity facilitate effective avoidance strategies at this age. During platform entries, BLA activity in adolescents ramped up prior to entry, while activity in juveniles only ramped up after they entered the safety platform (Fig. 2h), suggesting that distinct BLA activity patterns may underlie similar levels of PMA in juveniles vs. adolescents.

In NAc, only juveniles increased activity following tone onset (Fig. 2i). In adolescents, activity following tone onset was positively correlated with the latency to enter the platform and negatively correlated with platform bout length (Fig. 2j), indicating that elevated levels of NAc activity oppose threat avoidance. No such relationships between NAc activity and behaviour were observed in the other age groups. During PMA retrieval, all ages ramped up activity during entries onto the safety platform, and then decreased activity once animals were on the safety platform (Fig. 2k). Our findings suggest that the distinct trajectories of PL, BLA and NAc development may jointly determine developmental changes in threat avoidance behaviour, with changes in PL and NAc playing particularly important roles in adolescence. Further, these results suggest that changes in top-down PL-BLA and PL-NAc circuit function may underlie developmental changes in threat avoidance behaviour.

### Developmental switches in PL-BLA and PL-NAc roles during threat avoidance

In adult rodents, activating PL-BLA projections promotes threat avoidance while activating PL-NAc projections reduces threat avoidance behavior, promoting exploratory and reward seeking behaviors instead^29^. Human fMRI studies indicate that PFC-amygdala functional connectivity switches from positive to negative between childhood and adolescence^67,68^. However, how and when developing top-down PL circuits causally influence behaviour remains unknown. To investigate this, we used optogenetics to activate or inhibit the PL-BLA and PL-NAc pathways during threat avoidance behavior in juvenile, adolescent or adult mice. We injected AAVs encoding an excitatory opsin, an inhibitory opsin or a fluorophore control into PL and implanted bilateral optic fibres above either BLA or NAc. Two weeks after the viral injection, we trained juvenile, adolescent or adult mice in PMA without optogenetic stimulation (Extended Data Fig. 5). Then, during the retrieval session, we paired tone presentations with blue laser pulses (473nm, 50 ms, 15 Hz) to activate PL axon terminals expressing ChR2 or yellow light (635 nm, continuous) to inhibit PL axon terminals expressing Jaws (Fig. 3a,b). To understand both how and when PL-BLA or PL-NAc activity influences behavior, we plotted the fraction of time mice spent on the platform in 5 second bins that spanned the 30 second conditioned tone (Fig. 3c–f).

**Figure 3.**
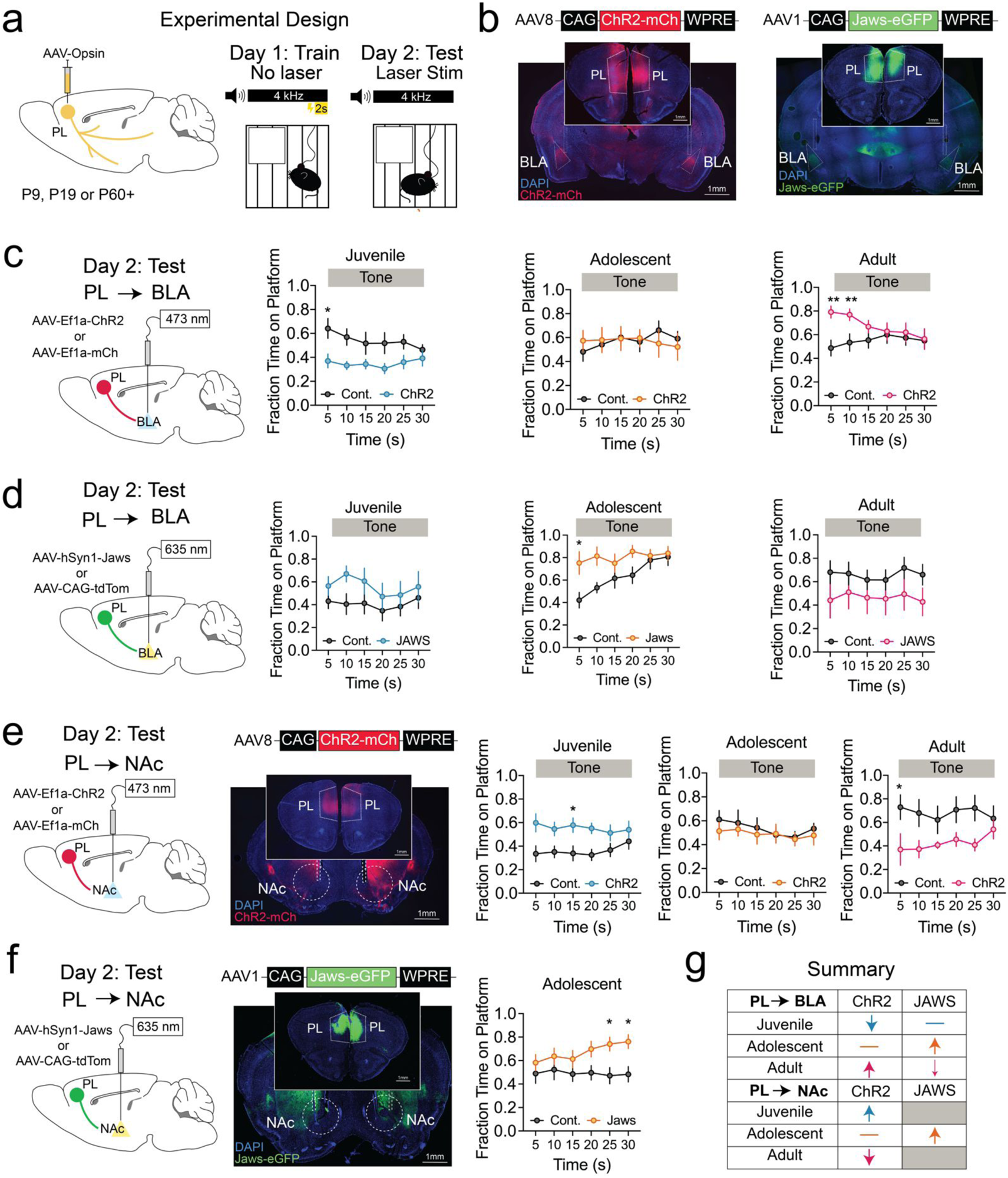
Manipulating PL-BLA and PL-NAc during PMA across development. **a,** Schematic of AAV injection into PL and PMA training and retrieval protocol. **b,** Representative image showing ChR2-mCherry and Jaws-GFP expression in PL neurons and axon terminals in BLA. **c,** Left: schematic of bilateral optic fiber placement over BLA and stimulation of PL terminals in this region during PMA retrieval; right: time on the platform across all ages, comparing performance of controls to ChR2 mice. Average time on platform is separated into 5 second bins across the 30 second tone period (Juvenile n=8 control, n=7 ChR2; Adolescent n=7 control, n=7 ChR2; Adult n=13 control, n=11 ChR2; 2-way ANOVA). **d,** Left: schematic of bilateral optic fiber placement over BLA and inhibition of PL terminals in this region during PMA retrieval; right: time on the platform across all ages, comparing performance of controls to Jaws mice (Juvenile n=9 control, n=10 Jaws; Adolescent n=9 control, n=8 Jaws; Adult n=9 control, n-9 Jaws; 2-way ANOVA). **e,** Left: schematic of bilateral optic fiber placement over NAc and stimulation of PL terminals in this region during PMA retrieval; right: time on the platform across all ages, comparing performance of controls to ChR2 mice (Juvenile n=10 control, n=6 ChR2, Adolescent n=6 control, n=8 ChR2, Adult n=6 control, n=7 ChR2; 2-way ANOVA). **f,** Left: schematic of bilateral optic fiber placement over NAc and inhibition of PL terminals in this region during PMA retrieval in adolescent mice. Right: time on platform across the tone period quantified, comparing Jaws mice to controls (Adolescent, n=10 control, n=11 Jaws; 2-way ANOVA). **g,** Summary of impact of PL-BLA and PL-NAc manipulations during PMA retrieval across all ages. Scale bars, 1 mm. Data represent mean ± s.e.m, *p<0.05, **p<0.01. Full statistical details can be found in Data Table 3.

We hypothesized that if weak PL control of subcortical structures leads to lower levels of threat avoidance in developing mice, then elevating activity in the PL-BLA pathway may increase PMA levels in juveniles and adolescents (Fig. 3c). Instead, we observed distinct effects in each age group. Activating the PL-BLA pathway decreased the amount of time juveniles spent on the platform, had no effect in adolescents, and consistent with previous studies, increased the amount of time adults spent on the safety platform, especially at the beginning of the tone (Fig. 3c).

To determine if these functional switches were causally related to behaviour, we optogenetically inhibited PL terminals in BLA during PMA retrieval. We found no significant effects on juvenile behavior. Surprisingly, the inhibiting PL-BLA activity significantly enhanced avoidance levels in adolescents. In adults, we observed a trend level decrease in time on the platform (Fig. 3d). We also analyzed the effects of PL-BLA inhibition on the latency to enter the platform after tone onset and on the number of platform entries during the tone. While we observed no significant effects in juveniles, adolescents had a trend-level decrease in latency to enter the platform and adults had a significant decrease in platform entries (Extended Data Fig. 6a–c). Taken together with the fibre photometry recordings, these data suggest PL may not play a causal role in threat avoidance in juveniles, but in adolescent and adults, the distinct levels of suppression following entry onto the safety platform – adolescents have less safety-induced suppression than adults (Fig. 2e) – is a key factor that determines the level of threat avoidance. In adolescents, relatively higher levels of PL-BLA activity while the animal is on the safety platform (Fig. 2e) may actually oppose threat avoidance, promoting exploration instead. In adults, a profound decrease in PL-BLA activity may keep mice on the safety platform, largely occluding the effects of optogenetic stimulation of PL-BLA.

We next tested whether changes in PL-NAc functions also contribute to developmental changes in behavior. Like what we observed with the PL-BLA pathway, activating PL-NAc produced opposing behavioural effects in juveniles compared to adults – opsin-expressing juveniles spent significantly more time on the platform than age-matched controls – and we observed no effect in adolescents. Consistent with previous studies, we found that in adults, activating PL-NAc projections decreased time spent on the platform (Fig. 3e). Inhibiting PL-NAc projections during presentations of the conditioned tone significantly increased the amount of time adolescent mice spent on the safety platform (Fig. 3f), indicating in adolescence, elevated activity in PL-NAc and PL-BLA both contribute to their lower levels of threat avoidance behaviour.

Given the differing effects of activating or inhibiting PL top-down circuits (Fig. 3g), we investigated whether activity in either pathway was innately appetitive or aversive. We placed mice in a real time place aversion assay in which one side of a chamber was paired with optogenetic excitation or inhibition of PL-BLA or PL-NAc terminals (Extended Data Fig. 7). We found no effects of exciting or inhibiting the PL-BLA pathway at any age (Extended Data Fig. 7a–f). Activating the PL-NAc pathway produced a place preference in adolescent mice but not in juveniles or adults (Extended Data Fig. 6g-l). Viral expression and fibre implants were confirmed via histology in all experiments (Extended Fig. 8a–h).

Taken together, these data indicate that when adolescents encounter threatening cues, elevated activity in both the PL-BLA and PL-NAc pathways promotes greater levels of exploration at the expense of threat avoidance. Further, our findings that activating the two pathways produced opposing behavioral effects in juveniles vs. adults suggest that the synaptic organization of these two pathways may undergo significant reorganization across those developmental transitions.

### Distinct trajectories of synaptic development in the PL-BLA and PL-NAc pathways

Synapse density and electrophysiological properties of neurons within mPFC undergo major changes throughout early life, including dendritic elaboration, growth and then pruning of dendritic spines, synaptic strengthening, and resting membrane potential hyperpolarization ^7,8,12,16,19,69^. The PL-BLA pathway undergoes a protracted maturation, with axons continuing to elaborate in BLA and synaptic excitation strengthening through adolescence^11^. However, how synaptic maturation progresses in the PL-NAc pathway, and how its trajectory aligns with maturation of the PL-BLA pathway, remains unknown. We examined synaptic density and synaptic strength in the PL-BLA and PL-NAc pathways across development to understand how they may jointly contribute to changes in threat avoidance behaviors.

We first set out to determine how PL axonal and synaptic density in BLA and NAc changes across the ages of interest. We injected into PL a viral vector designed to label putative presynaptic puncta (based on synaptophysin expression) and axons with bright green and red fluorophores, respectively (Fig. 4a). Allowing approximately two weeks for viral expression, we then perfused mice at P23, P35 or P60, immunostained brain sections, and imaged axons and putative presynaptic puncta in BLA and NAc using a confocal microscopy (Fig. 4b,c). To account for differences in the amount of viral transduction in PL, we normalized the axon density to the number of cell bodies we counted in PL. In NAc, PL axon density decreased significantly between adolescence and adulthood, but the bouton density (boutons/length of axon) remained constant across development (Fig. 4d). In BLA, PL axon density decreased significantly from the juvenile stage to adolescence and remained low into adulthood. We observed an increase in the bouton density in adults (Fig. 4e), suggesting that while many PL-BLA axons are pruned throughout adolescence, the remaining axons make additional synaptic contacts.

**Figure 4.**
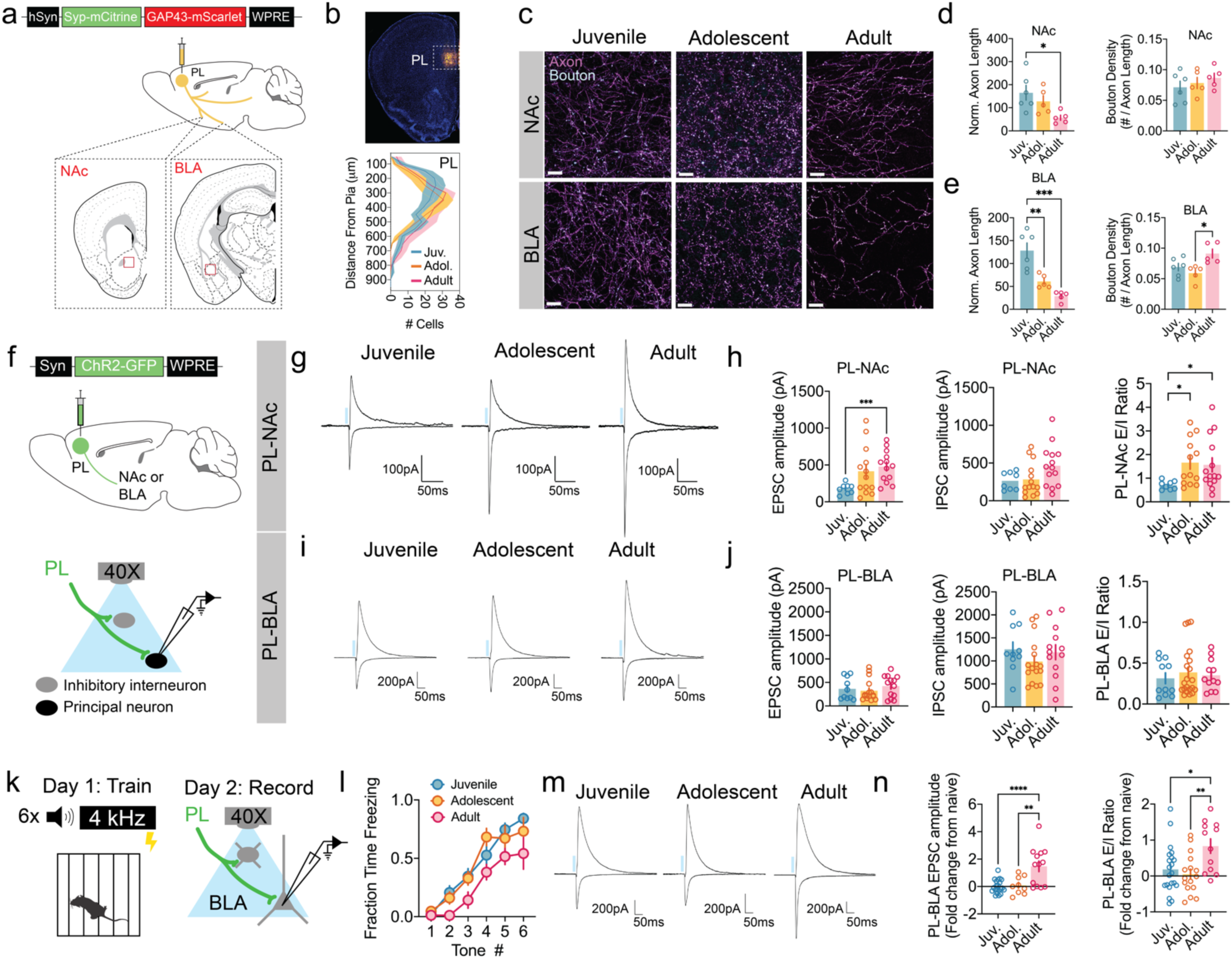
Synaptic Development of PL-NAc and PL-BLA. **a–e,** Anatomical interrogation of synaptic development of PL-NAc and PL-BLA (Juvenile n=6, Adolescent n=5, Adult n=5). **a,** Schematic of AAV injection into PL and regions of interest for confocal imaging. **b,** Example image of injection site and quantification of fluorescence across PL layers (2-way ANOVA). **c,** Representative images of axons and synaptophysin (syp) puncta in BLA and NAc at each age. **d,** Summary data of axon and bouton density in NAc in juvenile, adolescent and adult mice (One-way ANOVA). **e,** Summary data of axon and bouton density in BLA in juvenile, adolescent and adult mice (One-way ANOVA). **f–j,** Physiological interrogation of synaptic development of PL-NAc and PL-BLA in naïve mice (NAc: Juvenile n=8 cells from 5 mice, Adolescent n=14 cells from 4 mice, Adult n=14 cells from 4 mice; BLA: Juvenile n=11 cells from 5 mice, Adolescent 16 cells from 4 mice, Adult n=13 cells from 4 mice; One-way ANOVA). **f,** Schematic of AAV-ChR2 injection into PL and patch clamp recording configuration. **g,** Representative PL-evoked EPSCs (Vm=70mV) and IPSCs (Vm=0mV) recorded in NAc medium spiny neurons in juvenile, adolescent and adult mice. **h,** Summary data of PL-evoked synaptic currents in NAc across development. **i,** Representative PL-evoked EPSCs (Vm=70mV) and IPSCs (Vm=0mV) recorded in BLA pyramidal neurons in juvenile, adolescent and adult mice. **j,** Summary data of PL-evoked synaptic currents in BLA pyramidal cells across development. **k–n,** Physiological interrogation of synaptic development of PL-BLA in fear conditioned mice (Juvenile n=21 cells from 6 mice, Adolescent n=16 cells from 5 mice, Adult n=13 cells from 4 mice; One-way ANOVA). **k,** Schematic of fear conditioning followed by patch clamp recording configuration for recording PL-evoked synaptic currents in BLA. **l,** Summary data of freezing during fear conditioning. **n,** Summary data of PL-evoked synaptic currents in BLA pyramidal cells across development calculated as fold change from naive (trained/meannaive-1). Data represent mean ± s.e.m, *p<0.05, **p<0.01, ***p<0.001, ****p<0.0001. Full statistical details can be found in Data Table 4.

Next, we used channelrhodopsin-assisted circuit mapping^70^ to determine how the strength of synaptic transmission in PL-BLA and PL-NAc pathways changes over development. In naive mice, we injected AAV-ChR2 into PL. Allowing two weeks for viral expression, we then prepared acute brain slices from juvenile, adolescent or adult mice. We performed whole cell patch clamp recordings from neurons in BLA or NAc, recorded excitatory and inhibitory postsynaptic currents (EPSCs and IPSCs) evoked by optogenetically stimulated invading PL axon terminals (Fig. 4f). In the PL-NAc pathway, the amplitude of EPSCs increased steadily between juvenile and adult stages. On the other hand, the amplitude of IPSCs had a trend-level incrase between adolescence and adulthood. As a result, the ratio of excitatory to inhibitory currents (E/I ratio) in the PL-NAc pathway was significantly higher in adolescents and adults compared to juvenile mice (Fig. 4g,h). In the PL-BLA pathway, we observed modest increases in the amplitude of EPSCs and IPSCs from adolescence to adulthood, but no changes in the E/I ratio across those ages (Fig. 4i,j). These data suggest that different trajectories of synaptic development in the PL-NAc and PL-BLA pathways may contribute to differences in learned avoidance behaviour.

Previous studies showed that in adults, cued fear conditioning enhances the E/I ratio in the PL-BLA pathway^38^, suggesting that learning reorganizes top-down circuits. We performed a similar experiment to determine if the capacity for such plasticity changes over development. As described above, we injected AAV-ChR2 into PL. Two weeks later, when mice were juveniles, adolescents or adults, we performed auditory fear conditioning. This experimental design ensured that each mouse received the same number of footshocks, which can induce plasticity on their own. The next day, we prepared acute brain slices, performed whole cell patch clamp recordings from BLA principal neurons, and recorded EPCs and IPSCs evoked by optogenetic stimulation of PL axon terminals (Fig. 4k). Mice of all ages significantly increased their freezing levels during fear conditioning (Fig. 4l). However, we only observed changes in PL-BLA synaptic strength in adult mice. For each age group, we calculated fold-change in the strength of postsynaptic currents and the E/I ratio compared to naive controls (data shown in Fig. 5k). In adults, but not in juveniles or adolescents, we observed a significant fold increase in EPSC amplitude and in the E/I ratio (Fig. 4m,n), suggesting that adult-specific circuit-level plasticity in the PL-BLA pathway contributes to their elevated levels of PMA.

**Figure 5.**
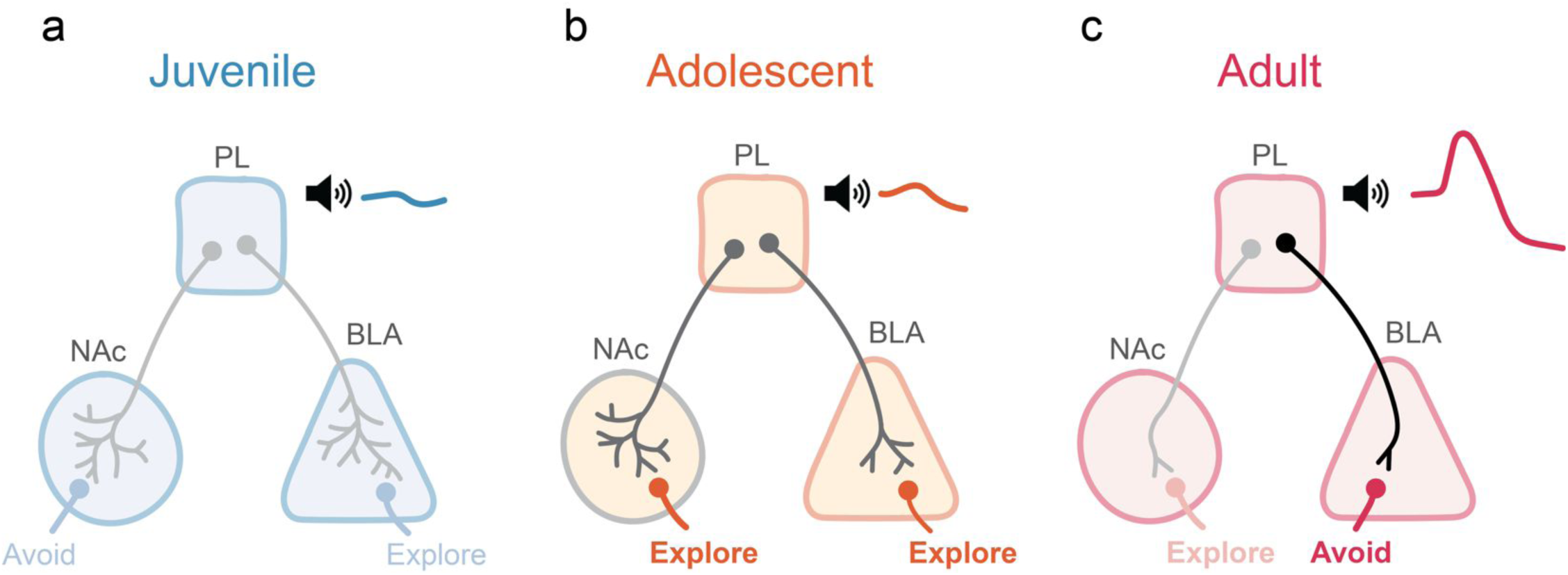
Proposed model of PL circuit organization contributing to threat avoidance. **a,** PL shows minimal dynamics to aspects of PMA. PL-BLA and PL-NAc circuits display a high degree of axonal arborization and are functionally wired to promote behavioral responses opposite to adults. Threat avoidance may require little prefrontal control in juveniles. **b,** In adolescence, both PL-BLA and PL-NAc circuits are wired to promote exploration at the expense of threat avoidance. During threatening cues, activity in these pathways reduces threat avoidance at this age. The PL-NAc pathway has increased in E/I ratio since the juvenile period while many PL axons have been pruned in BLA. **c,** In adults, axons in both PL-NAc and PL-BLA pathways have undergone significant pruning. Activity in the PL-BLA pathway promotes avoidance and has developed a capacity for learning-induced synaptic changes. The PL-NAc pathway is wired to reduce avoidance and is likely inhibited upon platform entry. PL displays robust modulation in response to threatening tones and threat avoidance behaviour.

## Discussion

We find that juvenile and adolescent mice display lower levels of threat avoidance compared to adults, each with unique behavioural signatures. Current models based on rodent and human evidence have suggested that weak prefrontal control over faster-developing subcortical limbic centers, leads younger animals to engage more often in impulsive or risky behaviors^22,23,48^. However, it remained unclear how progressive strengthening of top-down control can lead to nonlinear changes in behavior over development^24,25^. Here, for the first time, we established direct causal links between the maturation of frontolimbic circuits and developmental changes in threat avoidance behavior. Our studies reveal that pronounced rearrangements of frontolimbic circuit architecture, as opposed to linear strengthening with age, underlie developmental transitions in threat avoidance behavior.

In line with the idea of progressive increases top-down control, we observed increasingly robust PL modulation during threat cues and threat avoidance across juvenile, adolescent and adult stages^71^. However, functional manipulations revealed unique behavioural roles at each age.

In adults, we observed large increases in PL activity at the onset of a threatening cues followed by a strong suppression of activity when the animal reached safety. While adoelscents and juveniles had less pronounced mPFC modulation, PL activity was correlated with behaviour in adolescents but not in juveniles. Consistent with previous work^29^, we found that activity in the PL-BLA pathway promoted threat avoidance while activity in the PL-NAc pathway opposed it. Surprisingly, optogenetically activating PL-BLA and PL-NAc projections had opposite effects in juveniles compared to adults. We also found that the juvenile PL had made many synaptic contacts within BLA and NAc, that PL-NAc synapses were weaker than adults and that the PL-BLA pathway lacked adult forms of learning-induced plasticity. Together our data suggest that in the juvenile brain, promiscuous connectivity in top-down PL-circuits may promote competing behaviours when top-down pathways are active. Taken together with our recordings that show weak PL modulation during threat cues and the lack of effects of inhibiting juvenile PL-BLA during PMA, these data further suggest that PL has minimal influence over behaviour in juveniles.

Across species, adolescents engage in risky exploration^45,50,72^. Here we uncover a circuit mechanism that causally contributes to lower levels of threat avoidance in adolescence. We found that adolescents had lower threat cue-evoked activity compared to adults, but also less suppression of activity when animals entered the PMA platform, suggesting there may be more tonic PL activity in adolescents when they enter areas of safety. In line with this, inhibition of both the PL-BLA and PL-NAc pathways increased avoidance in adolescent mice. We observed significant pruning of the PL-BLA pathway between the juvenile stage and adolescence and only adults had learning-induced plasticity in the PL-BLA pathway. In the PL-NAc pathway, pruning occurred gradually between juvenile stages and adulthood, and synaptic excitation began to strengthen in adolescence, with a significantly higher E/I ratio than juveniles. These data suggest that an earlier strengthening of excitation in the PL-NAc pathway, combined with a still immmature PL-BLA pathway creates a circuit architecture in which, during encounters with threat, elevated PL activity promotes exploration at the expense of avoidance.

These findings refine previous models on developmental changes in threat behaviors^23,24,48^. While PL appears to become progressively more sensitive to threatening cues and safe locations with age, staggered trajectories in the synaptic maturation of PL-NAc and PL-BLA circuits underlie age-dependent switches in their behavioural roles across juvenile and adolescent stages.

Heterogeneous timelines in the maturation of top-down prefrontal circuits may be a key mechanism that shapes age-specific behaviours.

In humans and rodents, the juvenile and adolescent periods are sensitive windows when stressors can perturb brain development and threat-induced behaviors^66,73–76^. In humans, developmental and stress-induced changes in functional connectivity, region volume, and task-dependent dynamics have pointed to PFC, amygdala, and NAc as sensitive regions where external factors may alter activity, contributing to psychiatric diseases^66,67,77^. The lack of studies of the in vivo, causal functions mPFC, BLA and NAc in the developing brain have left a major gap in our understanding of how interactions between these regions produce developmental transitions in threat-induced behaviors. Our findings address this critical question, revealing how the staggered development of the PL-BLA and PL-NAc pathways establish distinct mPFC circuit architectures with unique behavioural roles. In revealing the processes by which top-down circuit maturation guides changes in threat-induced behaviors, we establish a foundation to understand how they can become disrupted.

## Methods

### Subjects

Female and male C57B16/J mice (JAX Stock No. 000664) were group housed (2–5 per cage). Infant mice were housed with their mothers and weaned at postnatal day 21. Mice were kept on a 12 hr light cycle (lights on 7am-7pm) in a temperature and humidity controlled room. Food and water were available ad libitum. All procedures followed animal care guidelines approved by the University of California, Los Angeles Chancellor’s Animal Research Committee.

### Behavioural Assays

#### Platform-mediated avoidance

Mice were handled for 3 days preceding the behavioural testing procedure. The conditioning chamber consisted of an 18 x 30 cm cage with a grid floor wired to a scrambled shock generator (Lafayette Instruments). The chamber was surrounded by a custom-built acoustic chamber and scented with 50% Windex. A thin acrylic platform (1.3 cm thick) covered 25% of the floor. Two small weigh boats filled with vanilla, almond or coconut extract were placed beneath the floor to encourage exploration of the chamber by the mice. During training on P23, P35 or in adulthood, mice were presented with three baseline 30s 4 kHz tones (CS), followed by nine presentations of the CS that co-terminated with a 2 s footshock (0.13mA).

The following day, mice were presented with six CS in the absence of shocks to probe ability to retrieve and express avoidance memory. Tones were separated by randomized interval lengths that ranged from 80 to 150 seconds. Non-shocked control mice underwent identical procedures except foot shocks were omitted.

#### Open-field Test

Mice were acclimated to the testing room for 10 minutes, and then placed in a plastic arena (50 x 50 x 40 cm). Locomotor activity and time spent in the center of the arena were recorded during a 10 minute test using the video-tracking system BioViewer. Using the tracking system, the arena was divided into two zones, the center (25% of the total area) and periphery (75% of the total area). Time spent in each zone as well as total distance traveled was recorded.

#### Shock Sensitivity

To assess the minimum foot-shock intensity required to elicit a behavioural response (vocalization, scurry or dart), naive P23, P35 and adult mice were placed in the same operant conditioning chamber as in PMA, but without the platform. Mice were exposed to a series of foot shocks, beginning at 0.02 mA and increased at 0.02 intervals until 0.20mA. The amplitude of the foot-shock at which a given mouse first vocalized, scurried and darted was recorded. Vocalization was defined as the emittance of an audible sound. Scurry was defined as rapid stepping with the absence of jumping. Dart was defined as a high velocity, horizontal jump.

#### Real-time place preference

To determine if optogenetic manipulation of mPFC-NAc or mPFC-BLA projections impacted behaviour beyond PMA, RTPP tests were performed the day following PMA retrieval for all optogenetic experiments. Following connection to the blue (stimulation; 473 nm; 15 Hz; SLOC Lasers, Shanghai, China) or yellow light (inhibition; 635 nm; 0 Hz; SLOC Lasers) laser, the mice were spaced in a place preference chamber (68 cm x 23 cm) for 20 min. For the first 10 min baseline period, mice were allowed to freely explore the apparatus and BioViewer software was used to track movement and determine which half of the chamber they preferred. This was used to determine which half of the chamber would receive laser stimulation or inhibition. For the following 10 min laser light was delivered on the preferred side of the chamber (PL-NAc inhibition and PL-BLA stimulation) or the non-preferred side (PL-NAc stimulation or PL-BLA inhibition). Results were calculated as percent change from baseline (test-baseline/baseline x 100).

### Optogenetic manipulation of mPFC-NAc and mPFC-BLA projections during PMA

#### Surgery

Juvenile cohorts (trained at P23) underwent viral injections at P9, adolescent cohorts (trained at P35) underwent viral injections at P19, and adult cohorts (trained or perfused after P60) underwent viral injections on or after P46. Mice were induced in 5% isoflurane in oxygen until loss of righting reflex and transferred to a stereotaxic apparatus. For P9 and P19 mice, the stereotax fitted with an attachment for developing mice including a small bite bar and developmental ear bars. A nonsteroidal anti-inflammatory agent was administered pre- and postoperatively to minimize pain and discomfort. The mouse’s head was shaved and prepped with three scrubs of alternating betadine and then 70% ethanol. Following a small skin incision, a dental drill was used to drill through the skull above the target. In P9 animals a 27-gauge syringe was used to poke a small hole in the skull. A syringe pump (Kopf, 693A) with a Hamilton syringe was used for pressure injections. For stimulation experiments, mice were bilaterally infused with adeno-associated virus (AAV) expressing either the excitatory opsin channelrhodopsin (AAV8-Ef1a-FAS-ChR2(H134)-mCherry-WPRE-pA, Addgene plasmid 37090, custom AAV produced from Janelia Virus Service). For inhibition experiments, mice were bilaterally infused with an AAV expressing the red-shifted inhibitory opsin Jaws (AAV5-hSyn-Jaws-KGC-GFP-ER2, Addgene #65014). Control mice were infused with a red fluorescent protein (excitation: AAV8-Ef1a-mCherry, Addgene #114470; inhibition: AAV1-CAG-tdTomato, Addgene #59462). Viral injection volume was scaled by age with 200nL injected at P9, 300nL injected at P19 and 700nL injected in adults. Virus was infused bilaterally at a rate 75nL/min into the mPFC (AP: +1.8; M: ±0.4; DV: -2.3mm). The syringe was left in the brain for 5-10 min following viral infusion in adult and adolescent mice. In juvenile mice the syringe was left in the brain for 2-5 minutes to decrease surgery time as mice were more sensitive to anesthesia. At P9 and P19, this coordinate was scaled based on the bregma-lamda distance using Neurostar StereoDrive Software (Neurostar GmbH, Tübingen, Germany). After recovery, P9 mice were returned to their home cage with their mother. All other ages the animals were housed with littermates.

Approximately 4 days prior to PMA training (10 days after viral infusion), juvenile and adolescent mice underwent an additional stereotaxic surgery to implant bilateral optical fibres. This was done to accommodate for skull growth during viral expression. Adult mice received fibre implants 2 weeks after viral infusion which was 1 week prior to PMA training. 200 um optical fibres (Braintech) were bilaterally implanted in the NAc (AP: +1.3; M: ±0.1; DV: -4.7mm) or BLA (AP: -1.6; M: ±3.0; DV: -4.3mm) and secured with Metabond (Parkell, NY, USA) to allow stimulation or inhibition of mPFC terminals in each of these areas.

#### Optogenetic manipulation during PMA

Prior to all experiments mice were habituated to optic tether (200 um, 0.22 NA, Doric Lenses, Quebec, Canada). No optogenetic manipulation occurred during PMA training. During training, mice that did not successfully avoid less than three shocks received an additional three tone shock pairings. For mice injected with ChR2, during the retrieval day, mPFC projections to either the NAc or BLA were photo-excited with a blue laser (473 nm; 15 Hz; SLOC Lasers) controlled by the behaviourDEPOT fear conditioning experimenter MATLAB app (Gabriel et al., 2022), during each of the six 30s tone presentations. The light power delivered, as measured through an optic fibre pre-implant, was set to output 10mW of light. For mice injected with Jaws, during the retrieval day, mPFC projections to either the NAc or BLA were photo-inhibited with constant yellow light laser (635 nm; SLOC Lasers). The light power delivered, as measured through an optic fibre pre-implant, was set to output 7.5mW of light.

### fibre photometry recordings during PMA

#### Surgery

Separate cohorts of each age were used to record bulk activity in PL, BLA or NAc. Mice were infused with AAV expressing the genetically encoded calcium indicator GCaMP6s (AAV9-GCaMP6s-WPRE-SV40, Addgene #100844). Virus (300 nL for all ages) was injected into either mPFC (AP: +1.8; M: ±0.4; DV: -2.3mm), NAc (AP: +1.3; M: ±0.1; DV: -4.7mm) or BLA (AP: -1.6; M: ±3.0; DV: -4.5mm). Infusions were done at a rate of 75nl/min and syringes left in place for 2-10 min, depending on the age. Similar to optogenetic experiments, optic fibre implants occurred in a separate surgery 4 days prior to PMA training. At this time the fibreoptic cannula (400um, 0.66 NA, Doric Lenses) was unilaterally implanted into the mPFC, NAc or BLA to allow for subsequent imaging of GCaMP fluorescence in cell bodies of each region.

#### Recordings during PMA

Fibre photometry was used to image bulk calcium activity in mPFC, NAc, and BLA neurons throughout PMA training and retrieval. Animals were habituated to the optical tether one day prior to recordings. During PMA training and retrieval we simultaneously imaged GCaMP6s and control fluorescence in each region using a commercial fibre photometry system and companion Synapse software controlling an RZ10x lock-in amplifier (Tucker Davis Technologies Inc., Alachua, FL). Two excitation wavelengths (465 and 405 nm) were modulated at 211 and 566Hz and filtered and combined by a fluorescence mini cube (Doric Lenses, Quebec, Canada). The combined excitation light was delivered via a 400 um core, 0.37 NA, low-fluorescence patch cord (Doric Lenses, Quebec, Canada) to an implanted optical fibre (fibre core diameter: 400 um; Doric Lenses). GCaMP6s emission fluorescence was collected through the mini cube and focused onto a femtowatt photoreceiver (Newport, <u>Model 2151, gain set to DC</u> <u>LOW</u>). LED power was set such that 80 and 20 units of light were received by the system for the 465 nm and 405 nm channel, respectively. Fluorescence was sampled at 1017 Hz, and demodulated by the processor. Time stamps for tone experiment start, finish and each tone onset were collected using TTLs sent from a custom MATLAB experiment designer (MathWorks, Natick, MA). Signals were saved using Synapse software and exported to MATLAB for analysis.

### Ex vivo electrophysiology

#### Surgery and Fear Conditioning

To optogenetically stimulate mPFC axons in the BLA and NAc, we injected 100nL of AAV8-Syn-ChR2(H134R)-GFP (Addgene #58880) into the right PL using techniques described above. Surgeries were performed 12-16 days prior to recording. For BLA recordings, naive and fear-conditioned animals were recorded on the same day using age- and sex-matched littermate controls. Fear conditioning was performed by delivering 6 20-second tones (4 kHz, 75 dB) co-terminating with a 2-second 0.5mA footshock. A randomized interval between 90 and 120 seconds separated each tone. Naive animals remained in their home cage. Recordings were performed the day after fear conditioning.

#### Acute Brain Slice Preparation

To prepare acute brain slices, mice were anesthetized with isoflurane and transcardially perfused with ice-cold slicing ACSF solution containing (in mM) 2.5 KCl, 1 NaH_2_PO_4_, 26.2 NaHCO_3_, 4 MgCl_2_, 11 Glucose, 210.3 Sucrose, 0.5 CaCl_2_, and 0.5 Na-Ascorbate (bubbled with 95% O_2_ / 5% CO_2_). The brain was rapidly dissected, and 300µm (BLA) or 230µm (NAc) sections were obtained from the hemisphere ipsilateral to the injection site using a Leica VT1200S vibratome. Slices were transferred to normal ACSF containing 125 NaCl, 2.5 KCl, 26.2 NaHCO_3_, 1 NaH_2_PO_4_, 2 MgCl_2_, 11 Glucose, and 2 CaCl_2_ (bubbled with 95% O_2_ / 5% CO_2_) and held at 34°C for 35-40 minutes. Slices were then allowed to cool to room temperature. Slices containing mPFC were also collected to verify the injection site.

#### Slice Electrophysiology

Recordings were performed at room temperature in normal ACSF. BLA and NAc were identified using white matter tracts and mPFC axon fluorescence. Cells were visualized under infrared-differential interference contrast through a 40x objective. Voltage clamp experiments were performed using borosilicate pipettes (5-7MΩ) filled with internal solution containing 117 CsMS, 20 HEPES, 0.4 EGTA, 2.8 NaCl, 5 TEA-Cl, 4 Na_2_-ATP, and 0.4 Na-GTP, adjusted to pH 7.3 using CsOH (280-290mOsm). In some NAc recordings, 1mM QX-314 was also included. Excitatory currents from mPFC terminal stimulation were obtained by holding neurons at -70mV and delivering 0.1ms (BLA) or 0.5ms (NAc) of 5-30mW blue (∼470nm) light through the 40x objective using a CoolLED pE-300Ultra light source. Inhibitory currents were recorded in the same way but holding neurons at 0mV.

Data were collected using a Multiclamp 700B amplifier and Digidata 1440A digitizer (Axon Instruments) with pClamp 10 (Molecular Devices). Recordings were sampled at 10kHz and filtered at 2kHz. Series resistance and input resistance were monitored throughout the experiment by measuring the capacitive transient and steady-state deflection in response to a 5mV test pulse, respectively. Series resistance was <25MΩ, did not change more than 20% throughout a session, and was not compensated. Data were analyzed in Python. Analysis was based on the average of 10 sweeps. Currents were analyzed relative to the baseline holding current. EPSCs and IPSCs were quantified by measuring the peak response when cells were voltage clamped at -70mV and 0mV, respectively. In some cases, a clear polysynaptic peak was present in the excitatory current after the initial monosynaptic peak. In these cases, the minimum value of the initial monosynaptic peak was used as the excitatory current. For comparisons of peak current values across conditions, analyses were restricted to cells with 15-30mW stimulation intensity. Statistical outliers were excluded via Grubbs’ test (alpha 0.01). The EPSC peak value was divided by the IPSC peak value for each cell to calculate the E/I ratio.

### Viral strategy to visualize synaptic innervation of NAc and BLA

#### Surgery

To visualize synaptic boutons from mPFC axons in the BLA and NAc we utilized a newly developed viral construct, (pAAV-hSyn-FLEx-loxP-Synaptophysin-mGreenLantern-T2A-GAP43-mScarlet, Schwarz Lab, St. Jude). AAV (serotype 2/9) was synthesized by the St. Jude Children’s Research Hospital Vector Development and Production Core to a final titer of 4.9×10^12. When in the presence of Cre, axons will express the red fluorescent protein mScarlet, and boutons the green fluorescent protein mGreenLantern. Stereotaxic surgeries were performed similarly to previous experiments, except the virus was delivered iontophoretically using glass micropipettes whose outside tip diameter measured 10-30um. A mixture of AAV-Syn-mGap-43-mScarlet and a AAV-Cre (AAV8-Ef1a-mCherry-IRES-Cre) was infused at a ratio of 3:1. A positive 5 uA, 7-second alternating injection current was delivered for 10 min (Stoelting Co.) to infuse the virus mixture and left in place for 2-10 min following infusion. The virus was allowed to express for 2 weeks and then brains were perfused for analysis.

### Histology

Following the behavioural experiments, mice were deeply anesthetized in a chamber filled with Isoflurane and transcardinally perfused with phosphate buffered saline (PBS), followed by 4% paraformaldehyde (PFA). Brains were removed and post-fixed in 4% PFA overnight, placed into 30% sucrose solution, then sectioned into 60um slices using a cryostat and stored in PBS or cryoprotectant. Images were acquired using Leica DM6 scanning microscope (Leica Microsystems, Germany) fitted with a 10x objective.

GFP immunofluorescence was used to confirm expression of GCaMP6s in cell bodies and Jaws expression in PL axons terminals. Floating coronal sections were washed 3 times in 1x PBS for 30 min and then blocked for 2hr at room temperature in a solution of 10% normal goat serum and 0.3% Triton X-100 dissolved in PBS. Sections were then washed 3 times in PBS for 15 min and incubated in 3% serum blocking solution containing chicken anti-GFP polyclonal antibody (1:2000; Aves Labs, Davis, CA) with gentle agitation at 4℃ overnight. Sections were next rinsed 3 times in PBS for 30 min and incubated with donkey anti-chicken IgY, AlexaFluor 488 conjugate (1:500; Jackson ImmunoResearch, West Grove, PA) in 0.5% serum blocking solution at room temperature for 2 hr. Sections were washed a final 2 times in PBS for 10 min.

RFP immunofluorescence was used to confirm expression of ChR2-mCherry, as well as mCherry and tdTomato control viruses in PL axon terminals. Floating coronal sections were washed 3 times in 1x PBS for 30 min and then blocked for 2hr at room temperature in a solution of 10% normal donkey serum and 0.3% Triton X-100 dissolved in PBS. Sections were then washed 3 times in PBS for 15 min and incubated in 3% serum blocking solution containing rabbit anti-RFP polyclonal antibody (1:2000; Rockland Immunochemicals, Pottstown, PA) with gentle agitation at 4℃ overnight. Sections were next rinsed 3 times in PBS for 30 min and incubated with donkey anti-rabbit IgY, Cyanine Cy3 conjugate (1:500; Jackson ImmunoResearch, West Grove, PA) in 0.5% serum blocking solution at room temperature for 2 hr. Sections were washed a final 2 times in PBS for 10 min.

RFP and GFP immunofluorescence were used to amplify expression of GAP43-mScarlet and Synaptophysin-mGreenLantern for visualization. Staining procedures were as described above. Samples were imaged on a 63x on a Leica STELLARIS confocal microscope (Leica Microsystems, Germany). Confocal z-stacks were analyzed in three dimensions (3D) using Imaris software (Oxford Instruments). First, we rendered a 3D surface of the mScarlet+ axons and masked out any red or green fluorescence outside the surface. Then we trained a filament classifier to detect fluorescent axons and measured the total summed length of all axonal segments in the region of interest. Next, we generated surfaces around the green fluorescent synaptic puncta. Surfaces were filtered based on the red fluorescence intensity – boutons were only counted if they were located within top 50% brightest axon fluorescence intensity.

### Data analysis

#### Behavioural analysis

High resolution videos of PMA were collected at 50 Hz using Chameleon3 USB cameras (Teledyne FLIR) Point tracking of videos were performed in DeepLabCut^78^ and behaviour analyzed using BehaviorDEPOT^79^. Custom MATLAB code was used to quantify time on platform, latency to enter the platform, freezing,

#### Fibre Photometry

Data were pre-processed using a custom-written pipeline in MATLAB. Prior to analysis the signal was downsampled by 10x. Using the polyfit function, the isosbestic signal was fit to the 465nm signal and this curve was subtracted from the 465nm channel. To align fibre photometry and behavioural data a lookup table was generated using linear interpolation between each TTL pulse to identify which behaviour frame lines up with each photometry frame. Z scores were calculated using a baseline period of -5 to 0 seconds relative to tone onset, and -20 to -15 seconds relative to platform entries. The average of all traces for an individual animal was calculated and used for analysis. To generate plots, each animal’s average trace was smoothed by averaging values from every 0.5 seconds and the mean ±SEM of smoothed traces across animals was displayed. Area under the curve (AUC) values were calculated from the average trace for each individual animal and peak values determine using the maximum Z-score during trace of interest.

### Statistical analysis

All statistical tests were performed in GraphPad Prism.

## Data and code availability

Custom MATLAB code available upon request.

## Acknowledgements

We thank the DeNardo and Wilke laboratories for project discussion; S. Wilke, P. Golshani, C. Portera-Cailliau, W. Hong, and A. Adhikari for comments on the manuscript. We acknowledge the Broad Stem Cell Research Center Microsopy Core at UCLA for providing access to microscopes and Imaris software. This work was supported by R01MH127214-01A1, a Klingenstein Fellowship in Neuroscience, and a Whitehall Research Grant to LAD, an NSERC Postgraduate Fellowship to CBK, an NSF GRFP to CMG and a T32NS048004 to MWG.

## Author Information Authors and Affiliations

University of California, Los Angeles, Los Angeles, CA, USA

Cassandra Klune, Caitlin Goodpaster, Michael Gongwer, Rita Chen, Nico Jones and Laura DeNardo

St. Jude Children’s Research Hospital, Memphis, TN, USA Lindsay Schwarz

## Contributions

CBK, CMG, MWG: conceptualization, data collection, data analysis, visualization, manuscript writing and editing. RC, NSJ: Data collection and analysis. CJG: Custom MATLAB code and data analysis. LAS: AAV cloning and production, manuscript editing. LAD: conceptualization, data collection, data analysis, visualization, manuscript writing and editing, supervision, funding acquisition, projection administration.

## Corresponding Author

Correspondence to Laura A. DeNardo.

## Ethics Declaration Competing Interests

The authors declare no competing interests.

