## Extended Data Figure 1 for "Developmentally distinct architectures in top-down circuits"

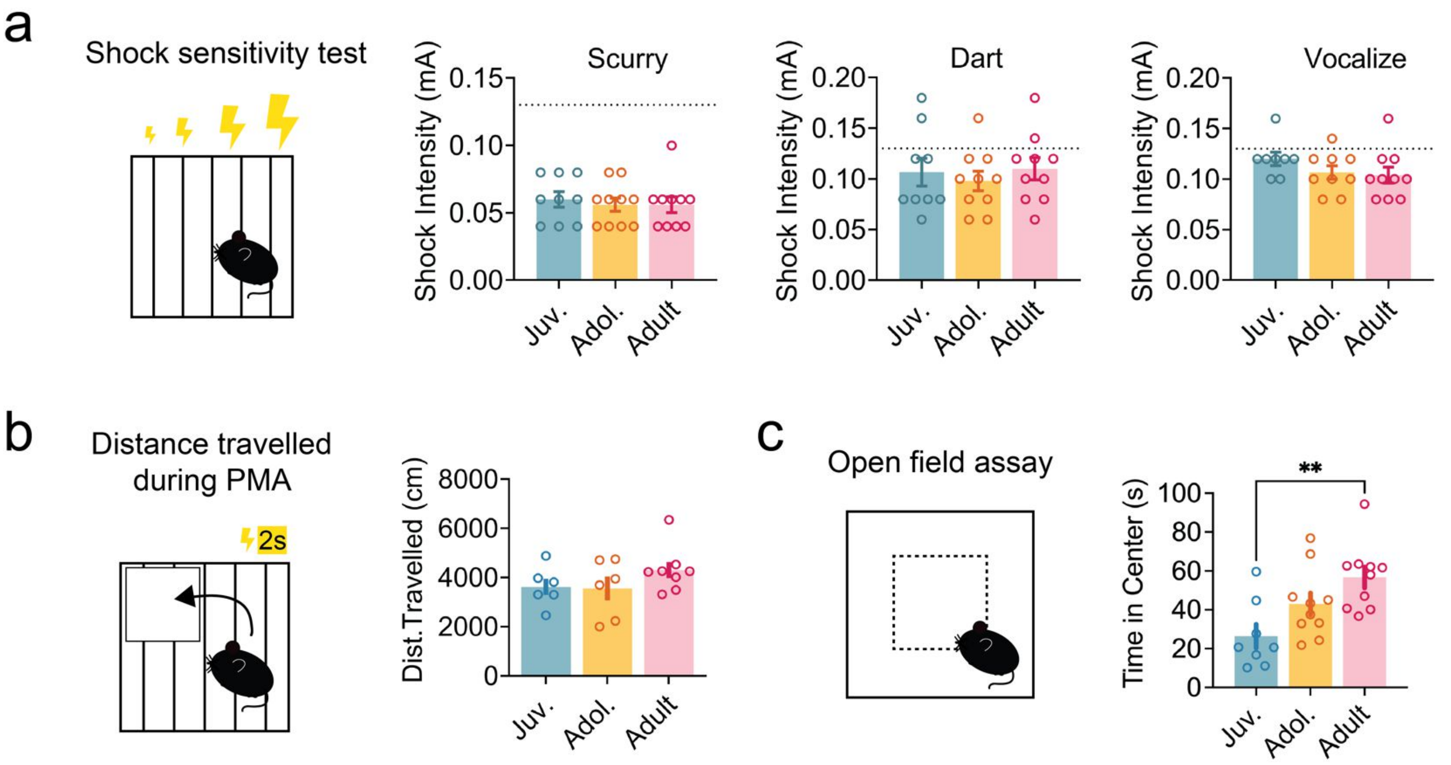


**Extended Data Figure 1. Shock thresholds in juvenile, adolescent and adult mice.**

**a,** Summary data of shock levels required to elicit behavioural repsonses to progressive increases in shock intensity (Juvenile: n=9, Adolescent n=10, Adult n=10; One-way ANOVA).

**b,** Summary data of distance traveled during PMA training (Juvenile n=6, Adolescent, n=6, Adult n=8; One-way ANOVA).

**c,** Summary data of time in the center of an open field (Juvenile n=8, Adolescent n=10, Adult n=10; One-way ANOVA).

Data represent mean ± s.e.m., **p<0.01. Full statistical details can be found in Data Table 1.
