## Extended Data Figure 2 for "Developmentally distinct architectures in top-down circuits"

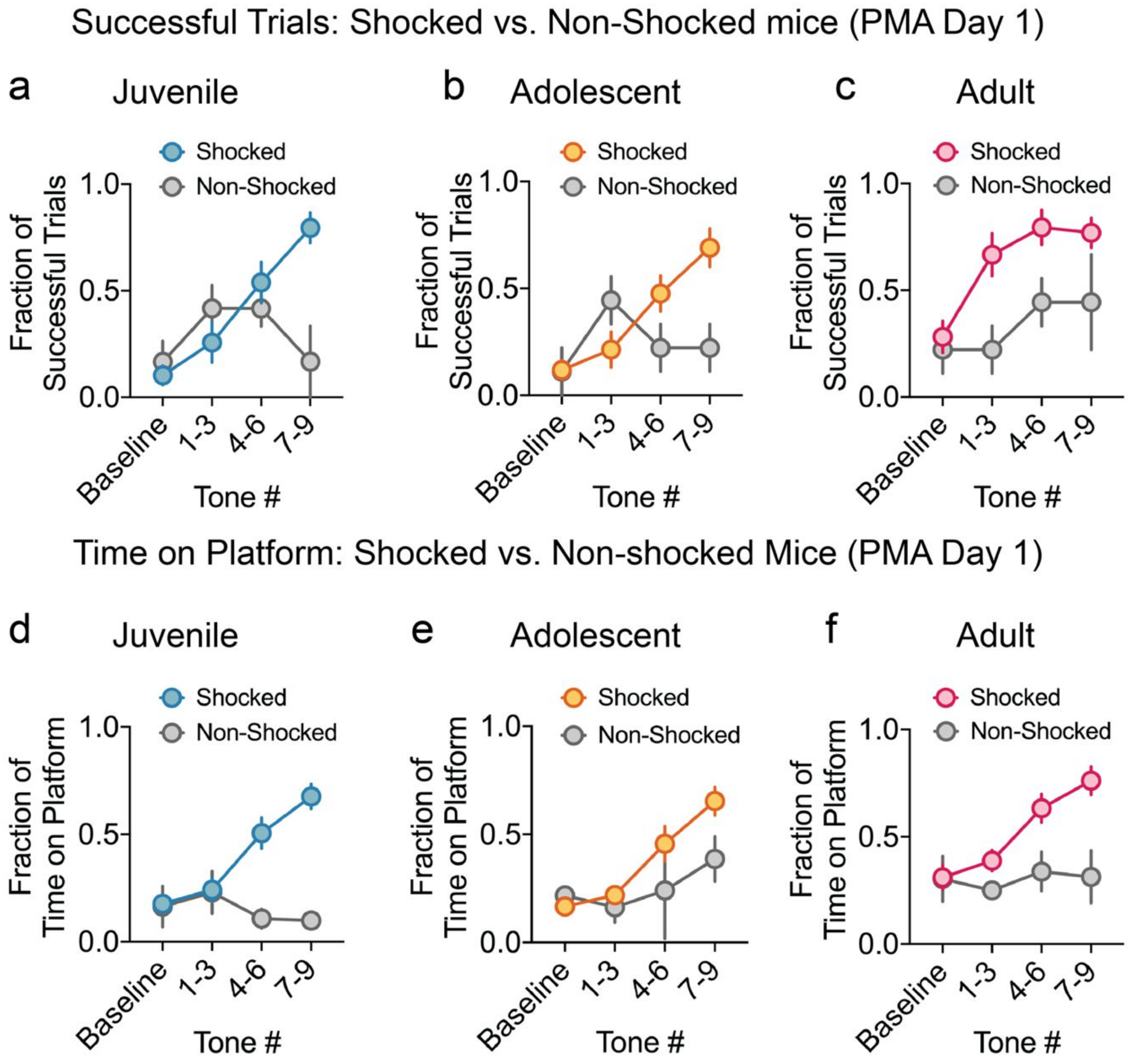


**Extended Data Figure 2. Behaviour in non-shocked control mice**

**a–c,** Comparing successful trials during PMA Day 1 for shocked vs. non-shocked (NS) control mice.
**a,** Summary data of successful trials for juvenile mice (n=4 NS, n=13 shocked; 2-way ANOVA).
**b,** Summary data of successful trials for adolescent mice (n=3 NS, n=14 shocked; 2-way ANOVA).
**c,** Summary data of successful trials for adult mice (n=5 NS, n=13 shocked; 2-way ANOVA).
**d–f,** Comparing time on platform during PMA Day 1 for shocked vs. non-shocked control mice
**d,** Summary data of time on platform for juvenile mice (n=4 NS, n=13 shocked; 2-way ANOVA).
**e,** Summary data of time on platform for adolescent mice (n=3 NS, n=14 shocked; 2-way ANOVA).
**f,** Summary data of time on platform for adult mice (n=5 NS, n=13 shocked; 2-way ANOVA).
Data represent mean ± s.e.m. Full statistical details can be found in Data Table 1.
