## Extended Data Figure 3 for "Developmentally distinct architectures in top-down circuits"

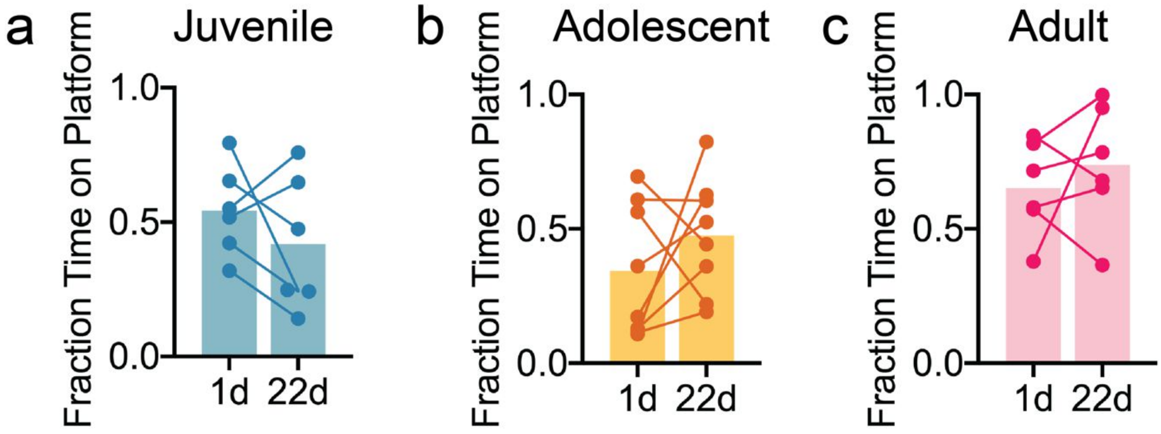


**Extended Data Figure 3. PMA behaviour during a second retrieval test.**
**a,** Fraction of time spent on platform during conditioned tone presentations during retrieval sessions occurring 1d or 22 days after training in (a) juvenile (n=6, *P*=0.31, Paired t-test), (b) adolescent (n=8, *P*=0.33, Paired t-test) and (c) adult mice (n=6, *P*=0.49, Paired t-test). Data represent mean ± s.e.m.
