## Extended Data Figure 4 for "Developmentally distinct architectures in top-down circuits"

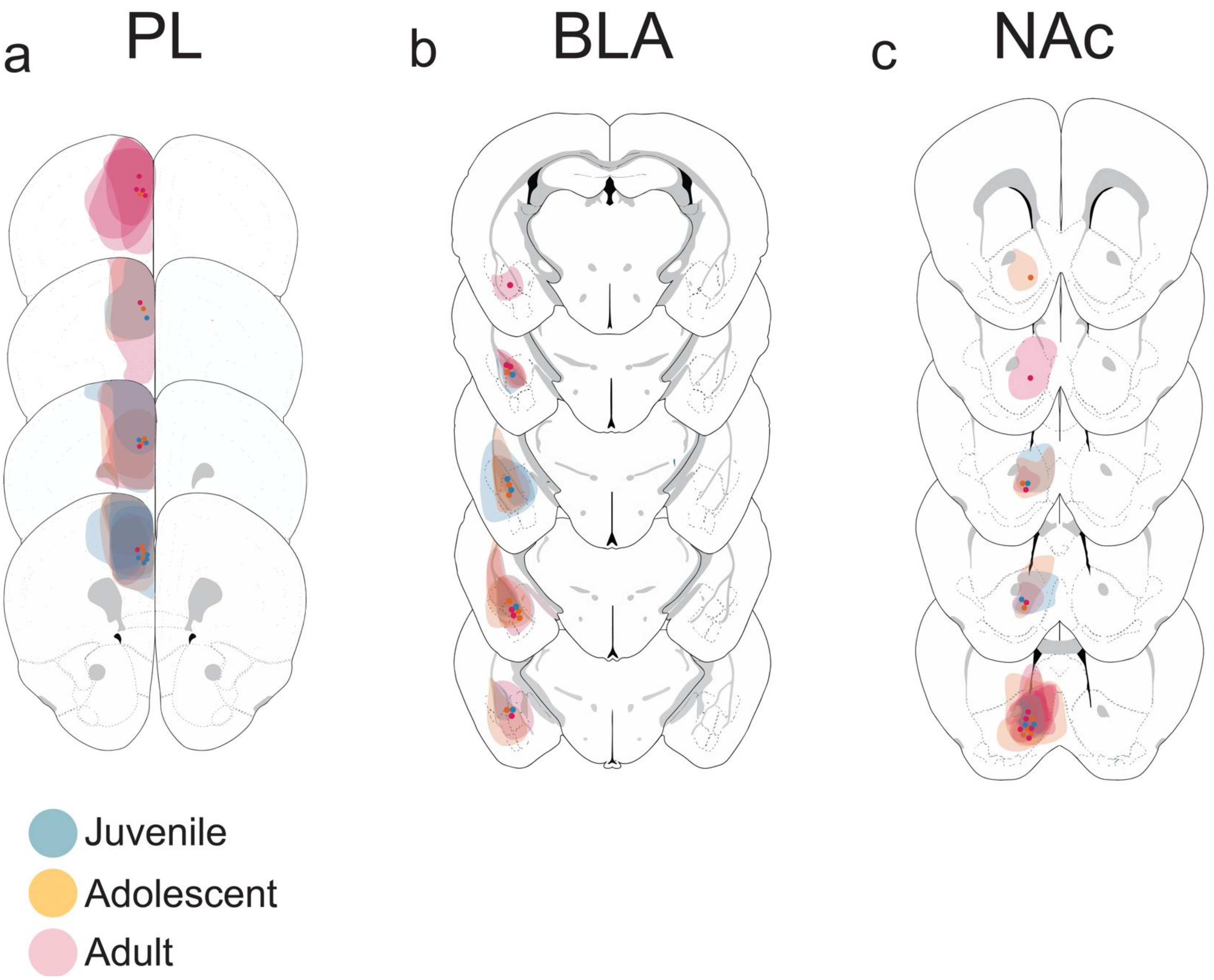


**Extended Data Figure 4: Histology maps for fiber placement in photometry studies**
**a,** Viral expression and fibre placement for all ages in PL.
**b,** Viral expression and fibre placement for all ages in the BLA.
**c,** Viral expression and fibre placement for all ages in the NAc.
