## Extended Data Figure 5 for "Developmentally distinct architectures in top-down circuits"

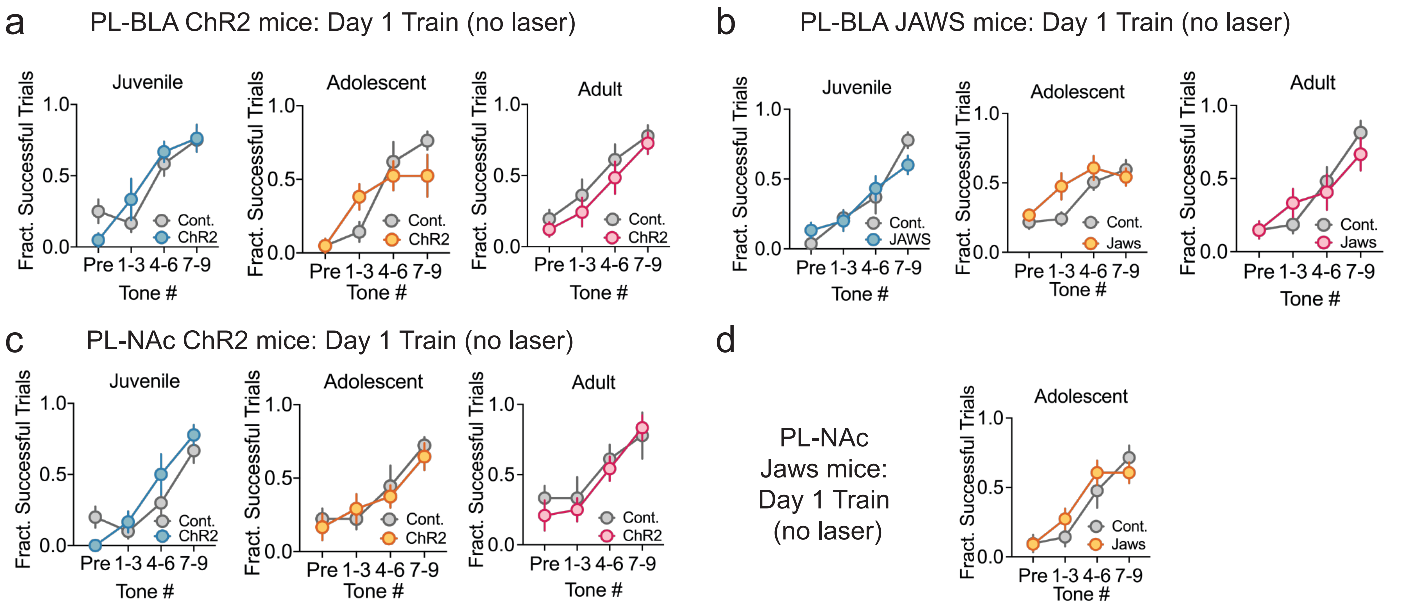


**Extended Data Figure 5. PMA training data for optogenetic manipulation experiments**

**a,** Fraction of successful trials during PMA training (laser off) for juvenile, adolescent for PL-BLA ChR2 stimulation experiments (Juvenile n=8 control, n=7 ChR2, Adolescent n=7 control, n=7 ChR2, Adult n=7 control, n=6 ChR2; 2-way ANOVA).

**b,** Fraction of successful trials during PMA training (laser off) for juvenile, adolescent for PL-NAc ChR2 stimulation experiments (Juvenile n=9 control, n=9 Jaws, Adolescent n=10 control, n=8 Jaws, Adult n=9 control, n=9 Jaws, 2-way ANOVA).

**c,** Fraction of successful trials during PMA training (laser off) for juvenile, adolescent for PL-BLA Jaws inhibition experiments (Juvenile n=6 control, n=6 ChR2, Adolescent n=7 control, n=7 ChR2, Adult n=6 control, n=7 ChR2; 2-way ANOVA).

**d,** Fraction of successful trials during PMA training (laser off) for juvenile, adolescent for PL-NAc Jaws inhibition experiment in adolescence (Adolescent n=11 control, n=12 Jaws).

Data represent mean ± s.e.m. Full statistical details can be found in Data Table 3.
