## Extended Data Figure 6 for "Developmentally distinct architectures in top-down circuits"

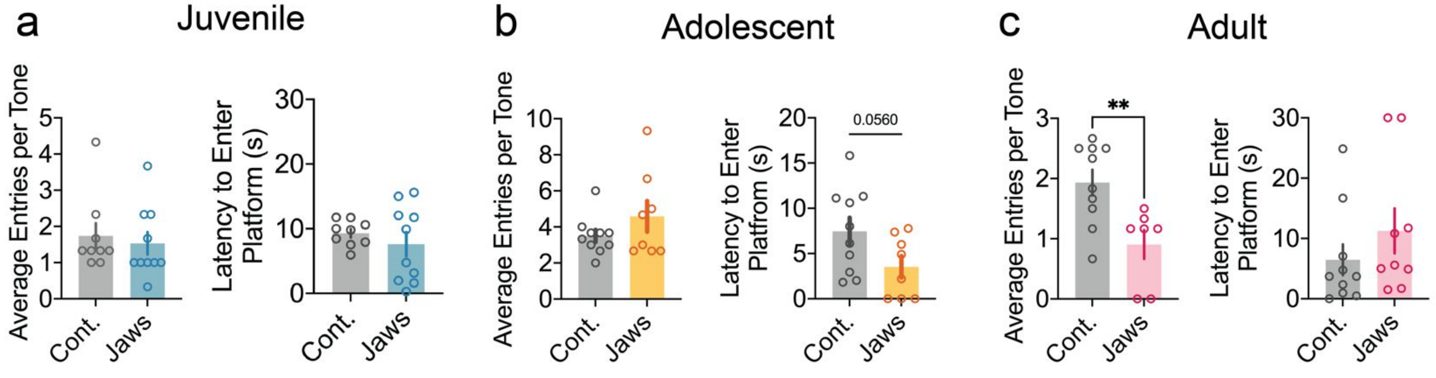


**Extended Data Figure 6. Behavioural effects of PL-BLA inhibition**
**a,** Platform entries (*P*=0.66, unpaired t-test) and latency to enter the platform (*P*=0.42, unpaired t-test) during PL-BLA inhibition in PMA retrieval in juveniles (n=9 control, n=10 Jaws).
**b,** Platform entries (*P*=0.27, unpaired t-test) and latency to enter the platform (*P*=0.05, unpaired t-test) during PL-BLA inhibition in PMA retrieval in adolescents (n=9 control, n=8 Jaws).
**c,** Platform entries (*P*=0.006, unpaired t-test) and latency to enter the platform (*P*=0.30, unpaired t-test) during PL-BLA inhibition in PMA retrieval in adults (n=9 control, n=9 Jaws).
Data represent mean ± s.e.m, **p<0.01.
