## Extended Data Figure 7 for "Developmentally distinct architectures in top-down circuits"

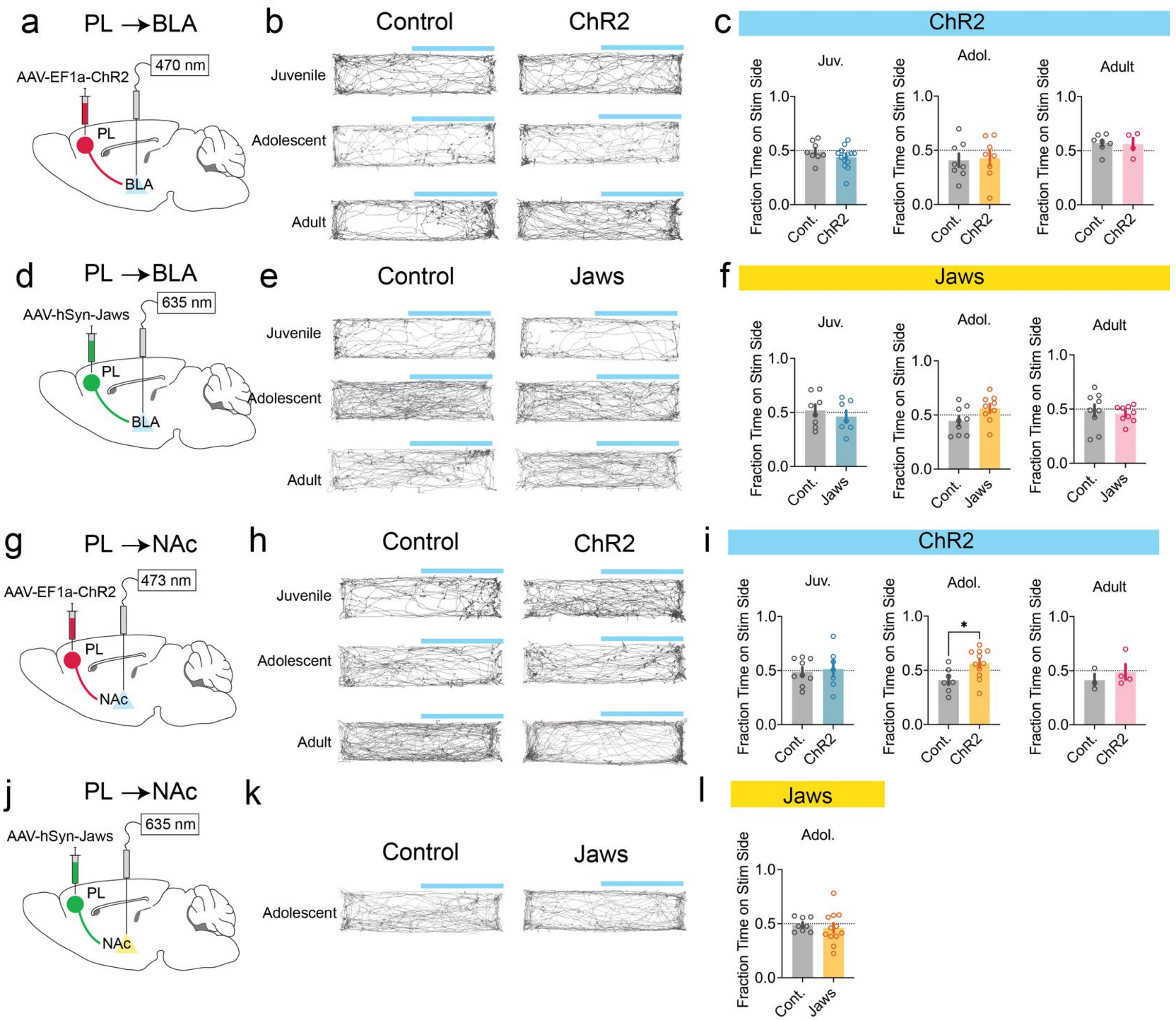


**Extended Data Figure 7: Real Time Place Preference (RTPP) for PL-BLA ChR2, PL-BLA Jaws and PL-NAc ChR2 and PL-NAc Jaws mice**

**a,** Schematic of AAV-ChR2 injection into PL and optic fibre implant over BLA.

**b,** Representative mouse trajectory maps during RTPP with PL-BLA stimulation for control and ChR2-expressing mice.

**c,** Summary data of fraction of time spent on the side of the chamber where laser stimulation occurred for PL-BLA ChR2 stimulation experiments (Juvenile: n=8 control, n=14 ChR2, Adolescent: n=8 control, n=8 ChR2, Adult: n=5 control, n=4 ChR2; Unpaired t-test).

**d,** Schematic of AAV-Jaws injection into PL and optic fibre implant over BLA.

**e,** Representative mouse trajectory maps during RTPP with PL-BLA inhibition for control and Jaws-expressing mice.

**f,** Summary data of fraction of time spent on the side of the chamber where laser stimulation occurred for PL-BLA Jaws inhibition experiments (Juvenile: n=9 control, n=7 Jaws, Adolescent: n=10 control, n=8 Jaws, Adult: n=9 control, n=9 Jaws; Unpaired t-test).

**g,** Schematic of AAV-ChR2 injection into PL and optic fibre implant over NAc.

**h,** Representative mouse trajectory maps during RTPP with PL-NAc stimulation for control and ChR2-expressing mice.

**i,** Summary data of fraction of time spent on the side of the chamber where laser stimulation occurred for PL-NAc ChR2 stimulation experiments (Juvenile: n=6 control, n=6 ChR2, Adolescent: n=7 control, n=7 ChR2, Adult: n=6 control, n=7 ChR2; Unpaired t-test).
**j,** Schematic of AAV- Jaws injection into PL and optic fibre implant over NAc.
**k,** Representative mouse trajectory maps during RTPP with PL-NAc stimulation for control and ChR2-expressing adolescent mice.
**l,** Summary data of fraction of time spent on the side of the chamber where laser stimulation occurred for PL-NAc ChR2 stimulation experiments (Adolescent: n=8 control, n=12 Jaws).
Data represent mean ± s.e.m. Full statistical details can be found in Data Table 3.
