## Extended Data Figure 8 for "Developmentally distinct architectures in top-down circuits"

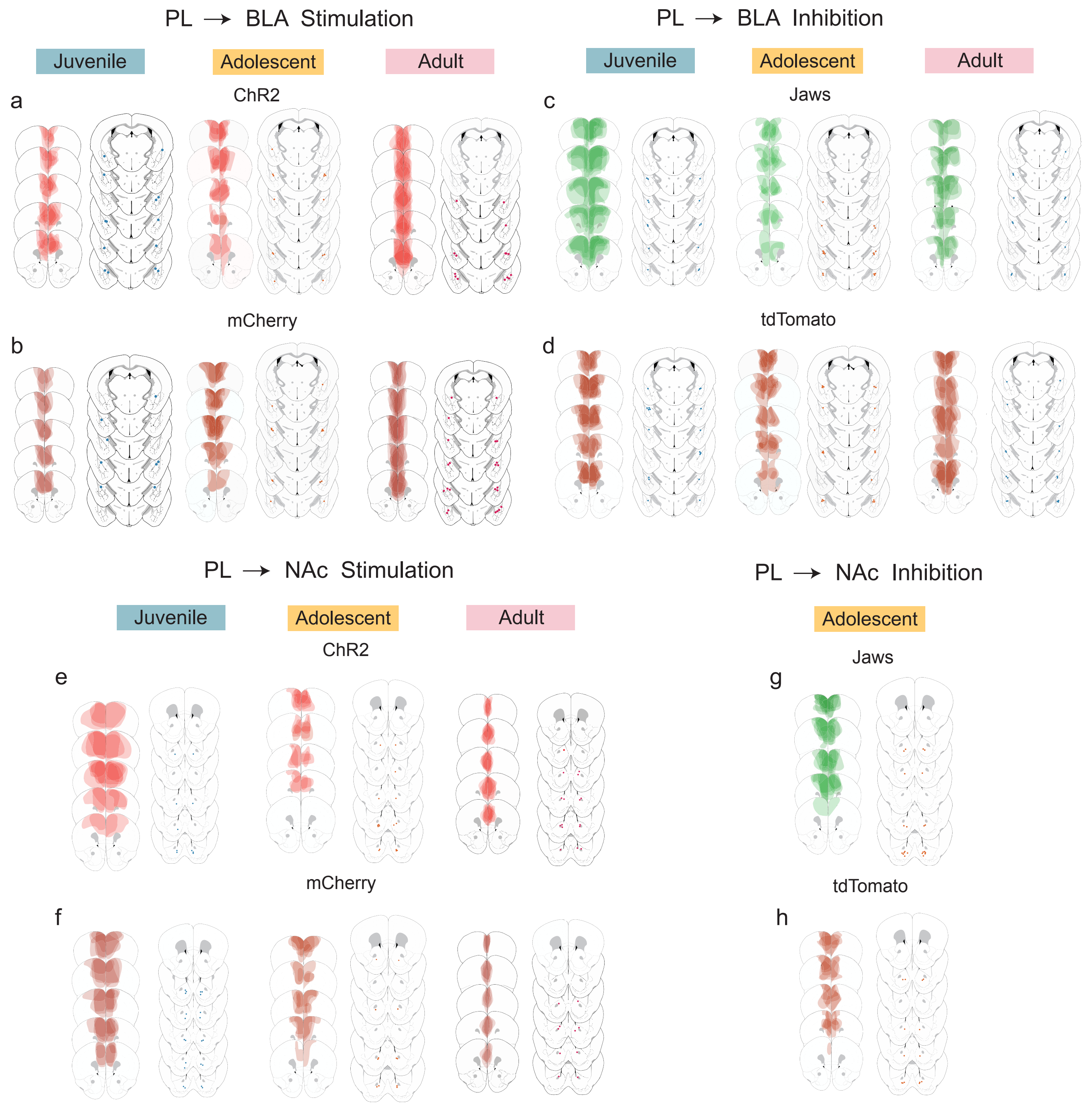


**Extended Data Figure 8: Histology maps for PL-BLA and PL-NAc manipulations.**
**a,** AAV-ChR2 viral expression in PL of juvenile, adolescent and adult mice and fiber implant placement above BLA.
**b,** AAV-mCherry control viral expression in PL of juvenile, adolescent and adult mice and fiber implant placement above BLA.
**c,** AAV-Jaws viral expression in PL of juvenile, adolescent and adult mice and fiber implant placement above BLA.
**d,** AAV-tdTomato control viral expression in PL of juvenile, adolescent and adult mice and fiber implant placement above BLA.
**e,** AAV-ChR2 viral expression in PL of juvenile, adolescent and adult mice and fiber implant placement above NAc.

**f,** AAV-mCherry control viral expression in PL of juvenile, adolescent and adult mice and fiber implant placement above NAc.
**g,** AAV-Jaws viral expression in PL of adolescent mice and fiber implant placement above NAc.
**h,** tdTomato control viral expression in PL of adolescent mice and fiber implant placement above NAc
