## Supplementary material for "Developmentally distinct architectures in top-down circuits": Data Table 1

**Data Table 1. Learned avoidance behaviour is developmentally regulated**

| | Two-way ANOVA with post hoc Tukey test | | | | | | | | | One-way ANOVA | | |
|---|---|---|---|---|---|---|---|---|---|---|---|---|
| **Figure 1** | $F_{(trial)}$ | DFn, DFd | p-value | $F_{(age)}$ | DFn, DFd | p-value | $F_{(interation)}$ | DFn, DFd | p-value | F | DFn, DFd | p-value |
| 1d: Sampe size | | | | | Juvenile n= 12, Adolescent n=13, Adult n=10 | | | | | | | |
| 1d. Train: Time on platform | 47.47 | 2, 76 | <0.0001 | 1.84 | 2,32 | 0.18 | 0.22 | 6, 96 | 0.97 | | | |
| 1d. Train: Successful trials | 28.98 | 2, 76 | <0.0001 | 4.82 | 2, 32 | 0.015 | 0.8944 | 6, 96 | 0.50 | | | |
| 1d. Train: Freezing | 34.89 | 2,60 | <0.0001 | 0.31 | 2, 32 | 0.74 | | | | | | |
| 1f-h: Sample size | | | | | Juvenile n= 12, Adolescent n=13, Adult n=10 | | | | | | | |
| 1f. Retrieval: Time on platform, All Trials | | | | | | | | | | 8.55 | 2, 32 | 0.001 |
| 1g. Retrieval: Time on platform, Binned Trials | 0.043 | 1,32 | 0.79 | 8.56 | 2, 32 | 0.001 | 4.31 | 2, 32 | 0.02 | | | |
| 1h. Retrieval: Freezing, Binned Trials | 9.84 | 1,21 | 0.004 | 0.26 | 2,32 | 0.77 | 1.28 | 2, 32 | 0.29 | | | |
| **Extended Data Figure 1.1** | | | | | | | | | | | | |
| 1.1a: Sample size | | | | | Juvenile n= 9, Adolescent n=10, Adult n=10 | | | | | | | |
| 1.1a. Scurry | | | | | | | | | | 0.17 | 2, 26 | 0.85 |
| 1.1a. Dart | | | | | | | | | | 0.31 | 2, 26 | 0.74 |
| 1.1a. Vocalize | | | | | | | | | | 1.37 | 2, 24 | 0.27 |
| 1.1b. Sample size | | | | | Juvenile n= 6, Adolescent n=6, Adult n=8 | | | | | | | |
| 1.1b. Distance traveled during PMA | | | | | | | | | | 1.29 | 2, 12 | 0.30 |
| 1.1c: Sample size | | | | | Juvenile n= 8, Adolescent n=10, Adult n=10 | | | | | | | |
| 1.1c. Time in Center of Open Field | | | | | | | | | | 6.74 | 2, 25 | 0.005 |

| | Two-way ANOVA with post hoc Sidak test | | | | | | | | | One-way ANOVA |
|---|---|---|---|---|---|---|---|---|---|---|
| **Extended Data Figure 1.2** | $F_{(trial)}$ | | | $F_{(shock)}$ | | | $F_{(int)}$ | | p-value | |
| 1.2a-f: Sample size | | | | | Juvenile n=4 non-shocked (NS), n=13 shocked; Adolescent n=3 NS, n=14 shocked; Adult n=5 NS, n=13 shocked | | | | | |
| 1.2a. Juvenile: Successful trial | 3.0 | 3, 45 | 0.04 | 1.5 | 1, 15 | 0.24 | 3.32 | 3, 45 | 0.028 | |
| 1.2b. Adolescent: Successful trial | 4.41 | 3, 45 | 0.008 | 2.51 | 1, 15 | 0.13 | 5.12 | 3, 45 | 0.004 | |
| 1.2c. Adult: Successful trial | 7.25 | 3, 48 | 0.0004 | 7.65 | 1, 16 | 0.014 | 2.05 | 3, 48 | 0.12 | |
| 1.2d. Juvenile: Time on platform | 4.17 | 3, 45 | 0.012 | 11.16 | 1, 15 | 0.0045 | 9.65 | 3, 48 | <0.0001 | |
| 1.2e. Adolescent: Time on platform | 5.67 | 3, 45 | 0.002 | 2.46 | 1, 15 | 0.14 | 1.23 | 3, 45 | 0.31 | |
| 1.2f. Adult: Time on platform | 8.78 | 3, 48 | <0.0001 | 9.17 | 1, 16 | 0.008 | 4.93 | 3, 48 | 0.005 | |

ANOVA Table for data shown in Figure 1 and Extended Data Figures.
