## Supplementary material for "Developmentally distinct architectures in top-down circuits": Data Table 2

**Data Table 2. Neural dynamics underlying PMA in PL, BLA and NAc**

| | | | | | | | | | | | | | | | |
|---|---|---|---|---|---|---|---|---|---|---|---|---|---|---|---|
| | | | | | | Two-way ANOVA | | | | | | | One-way ANOVA | | |
| **Figure 2** | $F_{(trial)}$ | DFn, DFd | p-value | | $F_{(age)}$ | DFn, DFd | p-value | | $F_{(interation)}$ | DFn, DFd | p-value | | F | DFn, DFd | p-value |
| 2b: Sample size | | | | | | | Juvenile n=16 , Adolescent n=21 , Adult n=20 | | | | | | | | |
| 2b. Successful Trials | 126 | 1, 53 | <0.0001 | | 3.41 | 2, 53 | 0.04 | | 1.45 | 2, 53 | 0.24 | | | | |
| 2b. Time on Platform | | | | | | | | | | | | | 0.26 | 2, 54 | 0.0003 |
| 2b. Freezing | | | | | | | | | | | | | 3.0 | 2, 54 | 0.058 |
| 2c-e: Sample size | | | | | | | Juvenile n=7 , Adolescent n=7 , Adult n=6 | | | | | | | | |
| 2c. PL Tone onset AUC | | | | | | | | | | | | | 10.19 | 2, 17 | 0.0012 |
| 2c. PL Tone onset Peak | | | | | | | | | | | | | 0.45 | 2, 17 | 0.0038 |
| | | | | | | Linear Regression | | | | | | | | | |
| | | Juvenile | | | | Adolescent | | | | Adult | | | | | |
| | F | DFn, DFd | $R^2$ | p-value | F | DFn, DFd | $R^2$ | p-value | F | DFn, DFd | $R^2$ | p-value | | | |
| 2d. PL Onset AUC x Latency | 0.65 | 1, 5 | 0.12 | 0.47 | 1.27 | 1, 5 | 0.20 | 0.31 | 4.80 | 1, 4 | 0.55 | 0.09 | | | |
| 2d. PL Onset AUC x Bout Duration | 4.35 | 1, 5 | 0.47 | 0.09 | 8.43 | 1, 5 | 0.63 | 0.03 | 4.19 | 1, 4 | 0.51 | 0.11 | | | |
| 2d. PL Onset AUC x Successful trials | 0.28 | 1, 5 | 0.05 | 0.62 | 17.75 | 1, 5 | 0.78 | 0.008 | 67.08 | 1, 4 | 0.94 | 0.001 | | | |
| 2e. PL Platform entries Post1 | | | | | | | | | | | | | 0.62 | 2, 17 | 0.23 |
| 2e. PL Platform entries Post2 | | | | | | | | | | | | | 2.91 | 2, 17 | 0.001 |
| 2f-h: Sample size | | | | | | | Juvenile n=5, Adolescent n=7 , Adult n=6 | | | | | | | | |
| 2f. BLA Tone onset AUC | | | | | | | | | | | | | 3.09 | 2, 15 | 0.075 |
| 2f. BLA Tone onset Peak | | | | | | | | | | | | | 1.01 | 2, 15 | 0.35 |
| 2g. BLA Onset AUC x Latency | 0.002 | 1, 3 | 0.0005 | 0.97 | 2.46 | 1, 5 | 0.33 | 0.18 | 0.05 | 1, 4 | 0.01 | 0.83 | | | |
| 2g. BLA Onset AUC x Bout Duration | 1.69 | 1, 3 | 0.36 | 0.28 | 0.97 | 1, 5 | 0.16 | 0.37 | 22.25 | 1, 4 | 0.85 | 0.009 | | | |
| 2g. BLA Onset AUC x Successful trials | 1.73 | 1, 3 | 0.37 | 0.28 | 0.11 | 1, 5 | 0.02 | 0.76 | 12.21 | 1, 4 | 0.75 | 0.025 | | | |
| 2h. BLA Platform entries Pre | | | | | | | | | | | | | 0.34 | 2, 15 | 0.02 |
| 2h. BLA Platform entries Post | | | | | | | | | | | | | 0.22 | 2, 15 | 0.16 |
| 2j-k: Sample size | | | | | | | Juvenile n=4 , Adolescent n=7 , Adult n=7 | | | | | | | | |
| 2i. NAc Tone onset AUC | | | | | | | | | | | | | 2.21 | 2, 15 | 0.145 |
| 2i. NAc Toneo onst Peak | | | | | | | | | | | | | 0.33 | 2, 15 | 0.20 |
| 2j. NAc Onset AUC x Latency | 12.01 | 1, 2 | 0.88 | 0.074 | 7.034 | 1, 5 | 0.58 | 0.045 | 0.929 | 1, 5 | 0.16 | 0.38 | | | |
| 2j. NAc Onset AUC x Bout Duration | 4.59 | 1, 2 | 0.70 | 0.166 | 17.96 | 1, 5 | 0.78 | 0.0082 | 0.569 | 1, 5 | 0.10 | 0.48 | | | |
| 2j. Onset AUC x Successful trials | 0.190 | 1, 2 | 0.087 | 0.71 | 1.41 | 1, 5 | 0.22 | 0.289 | 0.589 | 1, 5 | 0.11 | 0.478 | | | |
| 2k. NAc Platform entries Pre | | | | | | | | | | | | | 0.55 | 2, 16 | 0.49 |
| 2k. NAc Platform entries Post | | | | | | | | | | | | | 2.02 | 2, 16 | 0.62 |

ANOVA Table for data shown in Figure 2.
