## Supplementary material for "Developmentally distinct architectures in top-down circuits": Data Table 3

**Data Table 3. Manipulating PL-BLA and PL-NAc during PMA across development**

| | Two-way ANOVA Two-way ANOVA with post hoc Sidak test | | | | | | | | | Unpaired t-test |
|---|---|---|---|---|---|---|---|---|---|---|
| **Figure 3** | $F_{(time)}$ | DFn, DFd | p-value | $F_{(opsin)}$ | DFn, DFd | p-value | $F_{(interation)}$ | DFn, DFd | p-value | p-value |
| 3c: Sample size | Juvenile n=8 control, n=7 ChR2; Adolescent n=7 control, n=7 ChR2; Adult n=13 control, n=11 ChR2 | | | | | | | | | |
| 3c. Juv. PL-BLA ChR2 | 1.12 | 5,65 | 0.36 | 6.57 | 1, 13 | 0.02 | 1.29 | 5, 65 | 0.28 | |
| 3c. Adol. PL-BLA ChR2 | 0.18 | 5, 60 | 0.97 | 0.07 | 1, 12 | 0.79 | 0.92 | 5, 60 | 0.47 | |
| 3c. Adult PL-BLA ChR2 | 1.22 | 5, 110 | 0.31 | 0.55 | 1, 22 | 0.14 | 5.02 | 5, 110 | 0.003 | |
| 3d: Sample size | Juvenile n=10 control, n=10 Jaws; Adolescent n=9 control, n=8 Jaws; Adult n=9 control, n=9 Jaws | | | | | | | | | |
| 3d. Juv. PL-BLA Jaws | 0.85 | 5, 75 | 0.52 | 2.48 | 1, 15 | 0.14 | 0.37 | 5, 75 | 0.87 | |
| 3d. Juv. PL-BLA Jaws entries | | | | | | | | | | 0.14 |
| 3d. Adol. PL-BLA Jaws | 4.99 | 5, 75 | 0.0005 | 5.03 | 1, 15 | 0.04 | 2.48 | 5, 75 | 0.04 | |
| 3d. Adol. PL-BLA Jaws entries | | | | | | | | | | 0.68 |
| 3d. Adult PL-BLA Jaws | 1.61 | 5, 80 | 0.17 | 1.61 | 1, 16 | 0.22 | 0.81 | 5, 80 | 0.54 | |
| 3d. Adult PL-BLA Jaws entries | | | | | | | | | | 0.02 |
| 3e: Sample size | Juvenile n=10 control, n=6 ChR2; Adolescent n=6 control, n=8 ChR2; Adult n=6 control, n=7 ChR2 | | | | | | | | | |
| 3e. Juv. PL-NAc ChR2 | 0.29 | 2, 30 | 0.76 | 9.47 | 1, 14 | 0.008 | 0.74 | 5, 70 | 0.59 | |
| 3e. Adol. PL- NAc ChR2 | 0.95 | 5, 60 | 0.46 | 0.41 | 1, 12 | 0.53 | 0.19 | 5, 60 | 0.96 | |
| 3e. Adult PL- NAc ChR2 | 0.38 | 5, 55 | 0.88 | 6.55 | 1, 21 | 0.03 | 1.06 | 5, 55 | 0.38 | |
| 3f: Sample size | Adolescent, n=10 control, n=11 Jaws | | | | | | | | | |
| 3f. Adol. PL-NAc Jaws | 1.57 | 5, 95 | 0.18 | 4.51 | 1, 19 | 0.04 | 2.3 | 5, 95 | 0.05 | |
| **Extended Data Figure 3.1** | | | | | | | | | | |
| 3.1a: Sample size | Juvenile n=8 control, n=7 ChR2; Adolescent n=7 control, n=7 ChR2; Adult n=11 control, n=11 ChR2 | | | | | | | | | |
| 3.1a. PL-BLA ChR2 Juvenile | 22.28 | 3, 39 | <0.0001 | 0.06 | 1, 13 | 0.82 | 1.64 | 3, 19 | 0.20 | |
| 3.1a. PL-BLA ChR2 Adolescent | 17.65 | 2, 29 | <0.0001 | 0.14 | 1, 12 | 0.72 | 2.3 | 3, 36 | 0.09 | |
| 3.1a. PL-BLA ChR2 Adult | 18.58 | 3, 60 | <0.0001 | 1.00 | 1, 20 | 0.33 | 0.10 | 3, 60 | 0.96 | |
| 3.1b: Sample size | Juvenile n=9 control, n=10 Jaws; Adolescent n=9 control, n=8 Jaws; Adult n=9 control, n=9 Jaws | | | | | | | | | |
| 3.1b. PL-BLA Jaws Juvenile | 26.47 | 3,51 | <0.0001 | 0.04 | 1, 17 | 0.84 | 1.43 | 3, 51 | 0.24 | |
| 3.1b. PL-BLA Jaws Adolescent | 15.13 | 3, 45 | <0.0001 | 2.0 | 1, 15 | 0.18 | 2.16 | 3, 45 | 0.11 | |
| 3.1b. PL-BLA Jaws Adult | 19.73 | 3, 48 | <0.0001 | 0.06 | 1, 16 | 0.80 | 1.18 | 3, 48 | 0.33 | |
| 3.1c: Sample size | Juvenile n=10 control, n=6 ChR2; Adolescent n=6 control, n=8 ChR2; Adult n=6 control, n=8 ChR2 | | | | | | | | | |
| 3.1c. PL-NAc ChR2 Juvenile | 20.11 | 3, 42 | <0.0001 | 0.06 | 1, 14 | 0.82 | 1.64 | 3, 42 | 0.20 | |
| 3.1c. PL-NAc ChR2 Adolescent | 13.94 | 3, 36 | <0.0001 | 0.17 | 1, 12 | 0.69 | 0.35 | 3, 36 | 0.79 | |
| 3.1c. PL-NAc ChR2 Adult | 9,37 | 3, 36 | 0.0001 | 0.97 | 1, 12 | 0.34 | 0.29 | 3, 36 | 0.89 | |
| 3.1d: Sample size | Adolescent, n=10 control, n=12 Jaws | | | | | | | | | |
| 3.1d. PL-NAc Jaws Adolescent | 33.19 | 3, 57 | <0.0001 | 2.02 | 1, 19 | 0.17 | 1.07 | 3, 57 | 0.37 | |
| **Extended Data Figure 3.2** | | | | | | | | | | |
| 3.2c: Sample size | Juvenile, n=8 control, n=14 ChR2; Adolescent, n=8 control, n=8 ChR2; Adult n=7 control, n=4 ChR2 | | | | | | | | | |
| 3.2c. PL-BLA ChR2 Juvenile | | | | | | | | | | 0.22 |
| 3.2c. PL-BLA ChR2 Adolescent | | | | | | | | | | 0.84 |
| 3.2c. PL-BLA ChR2 Adult | | | | | | | | | | 0.88 |
| 3.2f: Sample size | Juvenile, n=8 control, n=7 Jaws; Adolescent, m=9 control, n=9 Jaws; Adult, n=9 control, n=9 Jaws | | | | | | | | | |
| 3.2f. PL-BLA Jaws Juvenile | | | | | | | | | | 0.47 |
| 3.2f. PL-BLA Jaws Adolescent | | | | | | | | | | 0.07 |
| 3.2f. PL-BLA Jaws Adult | | | | | | | | | | 0.65 |
| 3.2i: Sample size | Juvenile, n=9 control, n=7 ChR2; Adolescent, n=7 control, n=11 ChR2; Adult n=3 control, n=4 ChR2 | | | | | | | | | |
| 3.2i. PL-NAc ChR2 Juvenile | | | | | | | | | | 0.75 |
| 3.2i. PL-NAc ChR2 Adolescent | | | | | | | | | | 0.02 |
| 3.2i. PL-NAc ChR2 Adult | | | | | | | | | | 0.45 |
| 3.2l: Sample size | Adolescent, n=7 control, n=11 Jaws | | | | | | | | | |
| 3.2l. PL-NAc Jaws Adolescent | | | | | | | | | | 0.62 |

ANOVA Table for data shown in Figure 3 and Extended Data Figures.
