## Supplementary material for "Developmentally distinct architectures in top-down circuits": Data Table 4

**Data Table 4. Synaptic Development of PL-NAc and PL-BLA**

| | Two-way ANOVA with post hoc Tukey test | | | | | | | | | One-way ANOVA | | |
|---|---|---|---|---|---|---|---|---|---|---|---|---|
| **Figure 4** | F | DFn, DFd | p-value | $F_{(age)}$ | DFn, DFd | p-value | $F_{(interaction)}$ | DFn, DFd | p-value | F | DFn, DFd | p-value |
| 4b-e: Sample size | Juvenile n=6, Adolescent n=5, Adult n=5 | | | | | | | | | | | |
| 4b. PL Layer Distribution | $F_{distance}$ =105.8 | 19, 28 | P<0.0001 | 1.43e-03 | 2, 280 | >0.99 | | | | | | |
| 4d. PL-NAc Axon Length | | | | | | | | | | 4.60 | 2, 13 | 0.31 |
| 4d. PL-NAc Bouton Density | | | | | | | | | | 0.71 | 2, 13 | 0.51 |
| 4e. PL-BLA Axon Length | | | | | | | | | | 19.09 | 2, 13 | 0.0001 |
| 4e. PL-BLA Bouton Density | | | | | | | | | | 5.96 | 2, 13 | 0.015 |
| 4h: Sample size | Juvenile n=8 cells from 5 mice, Adolescent n=14 cells from 4 mice, Adult n=14 cells from 4 mice | | | | | | | | | | | |
| 4h. PL-NAc EPSC | | | | | | | | | | 4.73 | 2, 22 | 0.03 |
| 4h. PL-NAc IPSC | | | | | | | | | | 2.92 | 2, 33 | 0.07 |
| 4h. PL-NAc E/I Ratio | | | | | | | | | | 3.76 | 2, 26 | 0.049 |
| 4j: Sample size | Juvenile n=11 cells from 5 mice, Adolescent n=16 cells from 4 mice, Adult n=13 cells from 4 mice | | | | | | | | | | | |
| 4j. PL-BLA EPSC | | | | | | | | | | 0.66 | 2, 36 | 0.52 |
| 4j. PL-BLA IPSC | | | | | | | | | | 0.94 | 2, 36 | 0.4 |
| 4j. PL-BLA E/I Ratio | | | | | | | | | | 0.29 | 2, 41 | 0.75 |
| 4l-n: Sample size | Juvenile n=21 cells from 6 mice, Adolescent n=16 cells from 5 mice, Adult n=13 cells from 4 mice | | | | | | | | | | | |
| 4l. Freezing | $F_{trial}$=54.4 | 5, 60 | <0.0001 | 4.70 | 2, 12 | 0.03 | 1.01 | 10, 60 | 0.45 | | | |
| 4n. PL-BLA fear conditioned EPSC | | | | | | | | | | 12.3 | 2, 40 | <0.0001 |
| 4n. PL-BLA fear conditioned E/I Ratio | | | | | | | | | | 5.66 | 2, 46 | 0.006 |

ANOVA Table for data shown in Figure 4.
